# Dicer loss in Müller glia leads to a defined sequence of pathological events beginning with cone dysfunction

**DOI:** 10.1101/2025.01.30.635744

**Authors:** Daniel Larbi, Alexander M. Rief, Seoyoung Kang, Shaoheng Chen, Khulan Batsuuri, Sabine Fuhrmann, Suresh Viswanathan, Stefanie G. Wohl

## Abstract

**Purpose:** The loss of Dicer in Müller glia (MG) results in severe photoreceptor degeneration as it occurs in retinitis pigmentosa or AMD. However, the sequence of events leading to this severe degenerative state is unknown. The aim of this study was to conduct a chronological functional and structural characterization of the pathological events in MG-specific Dicer-cKO mice *in vivo* and histologically.

**Methods:** To delete Dicer and mature microRNAs (miRNAs) in MG, two conditional Dicer1 knock-out mouse strains namely RlbpCre:Dicer-cKO_MG_ and GlastCre:Dicer-cKO_MG,_ were created. Optical coherence tomography (OCT), electroretinograms (ERGs) as well as histological analyses were conducted to investigate structural and functional changes up to six months after Dicer deletion.

**Results:** Dicer/miRNA loss in MG leads to 1) impairments of the external limiting membrane (ELM) – retinal pigment epithelium (RPE), 2) cone photoreceptor dysfunction and 3) retinal remodeling and functional loss of the inner retina, 1, 3 and 6 months after Dicer loss, respectively, in both strains. Furthermore, in the Rlbp:Dicer-cKO_MG_ strain, rod photoreceptor impairment was found 4 months after Dicer depletion (4) accompanied by alteration of RPE integrity (5).

**Conclusions:** MG Dicer loss in the adult mouse retina impacts cone function prior to any measurable changes in rod function, suggesting a pivotal role for MG Dicer and miRNAs in supporting cone health. A partially impaired RPE however seems to accelerate rod degeneration and overall degenerative events.

## Introduction

Retinal degenerative diseases including age-related macular degeneration (AMD) or retinitis pigmentosa (RP) cause vision impairment and eventually lead to blindness. While many retinal dystrophies can be linked to mutations in specific genes, non-genetic / epigenetic factors also contribute to disease progression ^1–6^.

microRNAs (miRNAs) belong to non-coding RNAs and act as epigenetic factors by repressing translation. They play a role in retinal disease progression as well as very likely in disease onset and manifestation since some of them are cell type-specific (for reviews, see ^2^). miRNAs require processing before they obtain their mature functional form, which occurs via special enzyme complexes, including the Dicer complex. Hence, deleting Dicer1 in retinal cells (photoreceptors or MG) or the RPE is a very efficient way to study the role of cell type-specific mature miRNAs ^7–10^.

MG are the primary glia in the retina and fulfill a variety of functions in the healthy retina including but not limited to 1) feeding neurons with lactate, 2) neurotransmitter recycling (via glutamine synthetase), 3) water ion homeostasis (via water pores and ion pumps), 4) structural stability and integrity via their long processes that span the entire tissue and form the internal/inner limiting membrane (ILM) towards the vitreous and external or outer limiting membrane (ELM/OLM) towards the RPE, 5) cone photopigment recycling and 6) the formation of the blood retinal barrier (BRB) by wrapping blood vessels ^11–17^ ^18^. Because MG have such a plethora of functions, it is not far-fetched to assume that MG impairment would result in devastating outcomes. Indeed, partial elimination of MG via toxins leads to activation of the remaining glia, disruption of the BRB, and photoreceptor loss resembling degenerative diseases such as AMD, diabetic retinopathy or Macular Telangiectasia 2 (Mactel2) ^19, 20^. This suggests that MG are not only responders to damage but potential initiators of retinal degeneration /retinal dystrophies.

We previously established an MG-specific Dicer knock-out mouse and showed that the loss of mature miRNAs in the glia leads to a slow-progressing phenotype that resembles retinitis pigmentosa several months after the manipulation ^9^. Intriguingly, MG miRNAs were reduced by about 80% as early as one month after Dicer deletion ^9, 21^. The overall retinal histology, however, appeared unaltered at this time. Only occasionally, the tissue seemed to have some enlarged areas (bulgy appearance), but whether a postmortem event caused this outcome was unclear. Furthermore, even 3 months after Dicer loss, the overall retina was histologically intact with only occasional alterations at the cellular level (MG displacement). Later, though, several substantial alterations were found, including a thin outer nuclear layer (ONL). How this pathophysiological phenotype developed, when it manifested, and whether the RPE contributed to the outcome was however not investigated and is not known.

Therefore, the aims of this study were 1) to investigate the impact of the loss of Dicer in adult MG with regard to onset, manifestation, and progression of pathophysiological events in the retina, and to 2) elaborate the contribution of the RPE to the pathophysiological phenotype.

We used two MG-specific Dicer knock-out strains, i.e. the aforementioned Rlbp-CreERT:Dicer-cKO_MG_ mouse ^9^ as well as a newly generated Glast-CreER:Dicer-cKO_MG_ and conducted spectral domain-optical coherence tomography (SD-OCT) and electroretinograms (ERG) coupled with histological analysis. We analyzed the mice early (one month after Dicer deletion), intermediate (3-4 months after Dicer deletion) and late stage (6 months after dicer deletion). We found that the first dysfunction concerns cone photoreceptors, followed by rod loss and functional loss of inner retinal neurons at a late stage. The degenerative events were more accelerated in the Rlbp-CreERT:Dicer-cKO_MG_ mouse that also showed partial RPE impairments.

## Methods

### Animals and Cre induction

All mice used in this study were housed at the State University of New York, College of Optometry, in accordance with the Institutional Animal Care and Use Committee approved protocols (IACUC) and to the ARVO Statement for the Use of Animals in Ophthalmic and Vision Research. The *Rlbp1-creERT2* strain was obtained from Dr. Edward Levine, Vanderbilt University ^22^. The GlastCreERT strain (Tg[Slc1a3-cre/ERT]1Nat, ID 012586, generated by Dr. Jeremy Nathans), the *R26-stop-flox-CAG-tdTomato* strain (Ai14, #007908), as well as the Dicer conditional knockout strain (*Dicer^f/f,^* #006001, generated by Brian Harfe, ^23^) were obtained from Jackson Laboratories to create the following wildtype (wt) and Dicer-cKO (cKO) strains: *Rlbp1CreER: stop^f/f^-tdTomato* (referred to as wt) and corresponding *Dicer^f/f:^ Rlbp1CreER: stop^f/f^-tdTomato* (referred to as RlbpCre-Dicer-cKO), as well as *Glast1CreER: stop^f/f^-tdTomato* (wt) and corresponding *Dicer^f/f^*: *Glast1CreER: stop^f/f^-tdTomato* (referred to as GlastCre-Dicer-cKO). Males and females were used. Genotyping was done using the primers listed in Supplementary Table S1. Tamoxifen (Sigma, St. Louis, MO) was administered intraperitoneally at 75 mg/kg in corn oil for four consecutive days, for Dicer deletion at P11-14, to initiate the recombination of the floxed alleles. Furthermore, S129 mice (Rlbp1CreERT background strain), tdTomato mice (C57Bl6 background strain), and Cre-negative mice were used as controls as well and compared to tamoxifen-treated wildtype mice to exclude the possibility of a treatment effect or any abnormality of the wildtype controls used.

### Retinal spectral-domain optical coherence tomography (SD-OCT) imaging

*In vivo* retinal structure was assessed using the Envisu R2200 SD-OCT device (Bioptigen, Durham, NC). Animals were anesthetized via intraperitoneal injection of 75-100 mg/kg Ketamine and 5-10 mg/kg Xylazine dissolved in sterile saline. Pupils were dilated with phenylephrine hydrochloride (2.5%) and tropicamide (0.5%). Corneas were kept lubricated during scans using methylcellulose. Rectangular and radial volume scans (1.4 x 1.4 mm, 1000 A-scans/B-scan X 15 frames/B-scan) were obtained while centered on the optic nerve head. For image analysis, retinal layers were manually segmented in a 9X9 plot in four regional quadrants (superior-inferior, nasal-temporal) and thicknesses were measured using the DiverRelease_2_4 software. Total retinal thickness was defined as the distance from the inner border of the retinal nerve fiber layer (NFL) and the outer border of the retinal pigment epithelium (RPE). All nuclear and plexiform layers were analyzed including nerve fiber layer /ganglion cell Layer (NFL/GCL), inner plexiform layer (IPL), inner nuclear layer (INL), outer plexiform layer and outer nuclear layer (ONL). The region containing the inner and outer segments of photoreceptors and the RPE could not be distinguished with high resolution. Therefore, the distance between the outer border of the external limiting membrane (ELM, also known as outer limiting membrane, OLM) to the outer border of the RPE was consolidated as ELM-RPE area as described previously ^24, 25^. Thickness measurements were conducted in the central areas (∼650 μm radius from the optic nerve [ON]). To create a timeline of thickness changes in different retinal layers, we averaged the thicknesses measured at various retinal eccentricities in both the superior-interior and nasal-temporal directions for each time point of analysis.

### Electroretinogram (ERG) recordings

Mice were dark-adapted overnight and anesthetized–via an intraperitoneal injection of 75-100 mg/kg Ketamine and 5-10 mg/kg Xylazine dissolved in sterile saline. Recordings were performed in the dark room using the Espion Electrodiagnostic rodent system (Diagnosys LLC, Lowell, MA, USA). Pupils were dilated with 1% tropicamide ophthalmic solution and the eyes were lubricated using 1% methylcellulose. Animals were kept on a heat pad during the recording to maintain constant body temperature. The Espion Electrodiagnostic system uses a gold wire electrode as a recording electrode, with needles inserted into the cheek and back as reference and ground electrodes respectively with a hand-held portable stimulator as the light source. Full field 5 ms flashes were used to elicit responses. ERGs were obtained with light intensities ranging from 0.001 to 64 cd.s/m^2^ (scotopic) and 0.06 to 17.5 cd.s/m^2^ (photopic). Scotopic and photopic ERG a-wave amplitudes were measured from the baseline of the waveform to the trough of the initial negative deflection and the b-wave amplitudes were measured from the trough of the initial negative deflection to the peak of the subsequent positive deflection. For an in-depth analysis of the scotopic b-wave, the Naka-Rushton equation was used, V(I) = Vmax (I^n^) / (I^n^ + K^n^), Equation 1. The intensity–response amplitudes were fitted with this generalized equation; “I” is the stimulus intensity. “V” is the amplitude at stimulus intensity I. “Vmax” is the maximum/saturated amplitude and an indirect measurement for responsiveness. “K” is the semi-saturation constant at represents the stimulus intensity I at which half of the saturated amplitude Vmax is reached, hence K is an indirect measurement for sensitivity. n is the slope of the fit curve and an indirect measurement of gain/heterogeneity ^26, 27^.

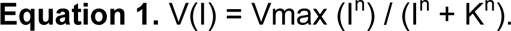

Generalized Naka-Rushton equation for estimating fit parameters.

### Tissue preparation

Mice were euthanized and eyes were marked at the nasal side, enucleated and transferred to pre-chilled phosphate-buffered saline (PBS) for eye cup preparation or Hank’s Balanced Salt Solution (HBSS) for RPE preparation.

#### Retinal cross sections

The eyeball was prefixed in 4% Paraformaldehyde (PFA, 4°C) for 20 min. After the removal of cornea, lens, iris, and vitreous body in PBS, eyecups were fixed in cold 4% PFA for additional 20 min, washed for 10 min in pre-chilled PBS, and incubated with 30% sucrose in PBS overnight at 4 °C. The tissue was embedded with nasal-temporal orientation in O.C.T. embedding medium and frozen at -80 °C. The frozen tissue was cross-sectioned in 12 μm thick sections for subsequent immunofluorescent labeling.

#### Retinal flat mounts

The unfixed eyeball was carefully dissected in cold PBS to remove cornea, lens, iris, ciliary body and vitreous body. The retina was carefully lifted from the RPE and transferred to a slide. To flatten the retina, 4 symmetric radial cuts were made. The tissue was subsequently fixed with cold 4% PFA for 20 min at room temperature (RT), washed three times for 10 min in PBS, and stained.

#### RPE flat mounts

RPE preparation was conducted as described before ^28^. In brief, muscles and connective tissue on the sclera were removed to facilitate the tissue flattening. Cornea, iris, lens and vitreous body were removed, and the RPE (with attached retina) carefully transferred on filter paper. Tissue was cut radially and the retina gently separated. The RPE (with sclera) was fixed with cold 4% PFA for up to 2 hours at RT, washed three times in PBS at RT, for 10 min and stained.

### Immunofluorescent labeling

For immunofluorescent staining, frozen sections were dried at 37°C for 20 min and fixed for 20 min with 4% PFA, and washed in PBS. Sections or flat mounts were incubated in blocking solution (5% milk block: 2.5 g nonfat milk powder in 50 mL PBS; with 0.5% Triton-X100, or 5% horse serum with 0.5% Triton-X100) for at least 1 hour at RT. After blocking, tissue was incubated with primary antibodies (Supplementary Table S2) in 5% milk block or horse serum overnight, retinal flat mounts were incubated for 2-3 days. After three thorough washes in PBS, tissue was incubated with secondary antibodies (Invitrogen/Molecular Probes, and Jackson ImmunoResearch, 1:500–1,000, Supplementary Table S2) for 1 hour (slides) or one day (flat mounts) at RT and counterstained with 4’,6-diamidino-2-phenylindole (DAPI, Sigma, 1:1,000). After incubation, tissue was washed three times in PBS and mounted with cover glasses and mounting medium (Invitrogen).

### Fluorescence microscopy, confocal laser scanning microscopy, and image processing

For whole tissue imaging, retinal eye cups were imaged using a fluorescent microscope (Keyence) equipped with a 4× objective. Immunofluorescent labeled retinal cross sections, RPE flat mounts and retinal flat mounts were imaged using a confocal laser scanning microscope (Olympus FM1200 Fluoview) equipped with 4×, 10×, 20× or 40× objectives and the Fluoview FV31S software. Images were processed and analyzed using Adobe Bridge, Adobe Photoshop, and Affinity.

### Statistical Analysis

Statistical analyses were performed using the Mann-Whitney U-test. Holm-Bonferroni method was used to correct for multiple comparisons. A p-value less than 0.05 was considered significant.

## Results

### The MG-specific Rlbp-Cre:Dicer-cKO_MG_ mouse is a reliable mouse model resulting in a robust phenotype resembling retinal degenerative diseases

In order to perform an *in vivo* in-depth analysis of retinal health and function in the MG-specific Dicer-cKO mouse model (referred to as Rlbp-Cre:Dicer-cKO_MG_) ^9^, we first validated the establishment of the reported phenotype. Tamoxifen was injected on postnatal day (P) 11-14 ^9^ to activate the Cre recombinase and the tissue was evaluated histologically one and 6 months after Dicer deletion (Figure 1A). The wildtype showed sufficient reporter induction (∼ 98%) in glutamine synthetase (GS) and Sox9 expressing MG (Figures 1B-G), consistent with previous reports ^9, 29^. The retinas of the 1-month old Rlbp-Cre:Dicer-cKO_MG_ showed complete reporter induction throughout the retina as well as the previously reported random areas with somewhat dislocated MG (Figures 1H-J). In the retinas of the 6-month old Dicer-cKO_MG_ mice, severe disruptions and thinning of the central areas were found, predominantly in the outer nuclear layer (ONL), confirming the reported phenotype^9^. The MG appeared to form a large glial seal (Figures 1K-L, arrowheads) resembling the events reported in late-stage retinal remodeling ^30^ and had reduced glutamine synthase labeling (Figure 1M arrows), a hallmark of specific gliosis ^30^. Furthermore, the RPE seemed to interact with the ELM/retina (Figure 1K, arrows), a process that appears to occur in a variety of retinal diseases with photoreceptor loss ^31, 32^. Hence, all previously reported features of this mouse model were present, confirming the robust and reliable phenotype and allowing an in-depth *in vivo* structural and functional analysis to elaborate the sequence of events leading to this severe degenerative state.

**Figure 1:**
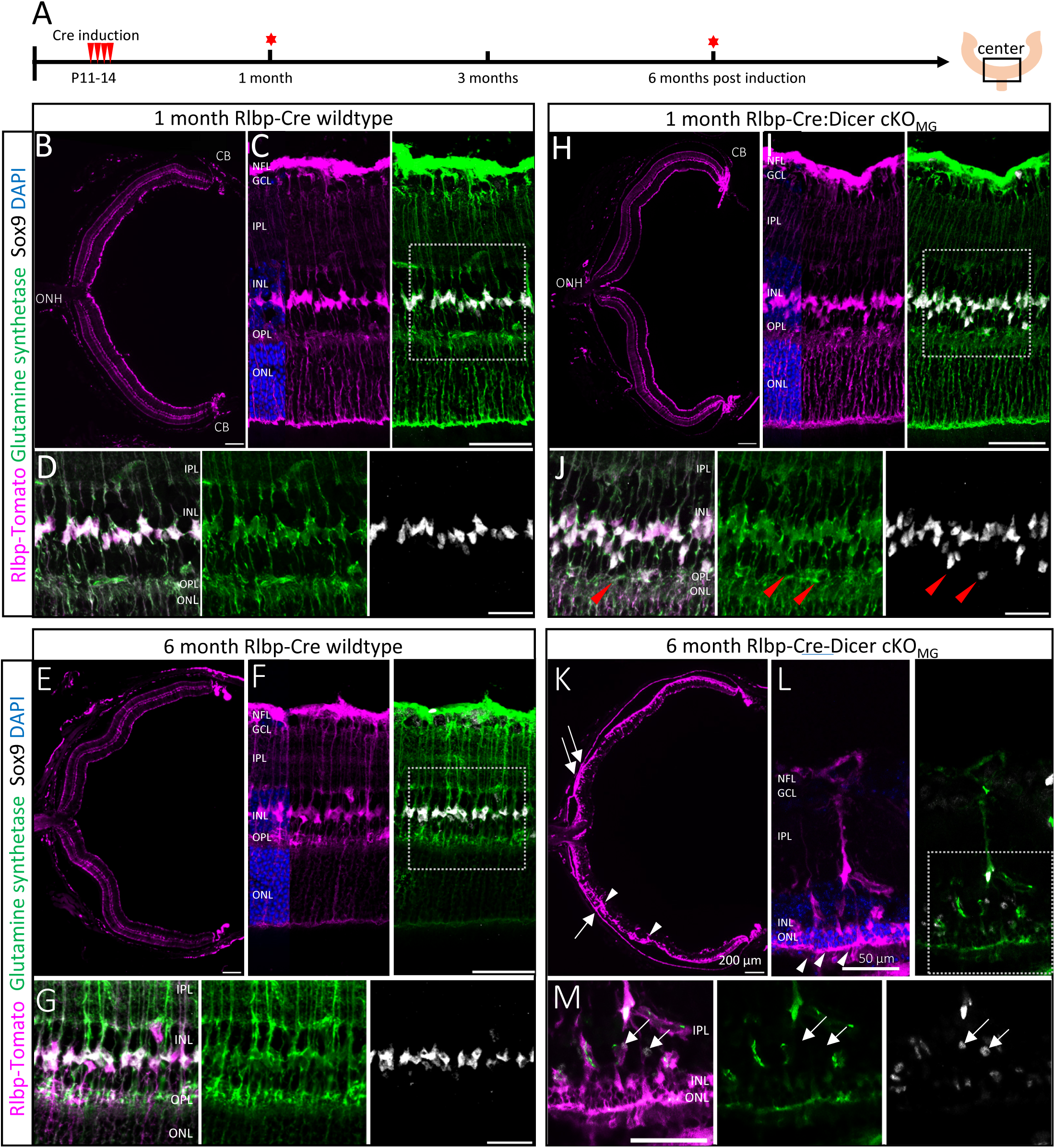
miRNA loss in Müller glia (MG) results in retinal degeneration at late stage. **A**: Experimental design showing Cre activation at P11-14 and time points of analysis one and six months after Cre induction (red stars). **B-M**: Immunofluorescent labeling with antibodies against RFP (Rlbp1:Tomato), Glutamine Synthetase (GS) and Sox9, as well as DAPI nuclear staining of center areas of wildtype (B-G) or Rlbp-Cre:Dicer cKO_MG_ mice (H-M), 1 or 6 months after Cre induction. The insets in C, F, I and L are shown in D, G, J and M respectively. Red arrowheads in J indicate displaced Müller glia (MG). Arrows in K show RPE-ELM interaction. Arrowheads in K and L show MG seal formation. Arrows in M show GS negative cells. Scale bars in B, E, H, K: 200 µm, in C, F, I, L: 50 µm, in D, G, J, M: 25 µm. NFL: Nerve Fiber Layer, GCL: ganglion cell layer, IPL: inner plexiform layer, INL: inner nuclear layer, OPL: outer plexiform layer, ONL: outer nuclear layer., ELM: external limiting membrane.

### miRNA loss in Müller glia does not result in substantial structural or functional retinal impairment in the early phase but leads to some alterations in the ELM-RPE and INL

To investigate structural abnormalities *in vivo*, we conducted spectral-domain optical coherence tomography (SD-OCT) one month after Dicer deletion, the time point by with ∼70-80% of all miRNAs are reduced ^9, 21^ (Figure 2A). Thickness measurements for whole retinal thickness as well as for the individual layers were conducted in the central areas (∼650 μm radius from the optic nerve [ON], across the nasal-temporal as well as the superior-inferior axis (Figures 2D, Supplementary Figures S1A, S2A)).

**Figure 2:**
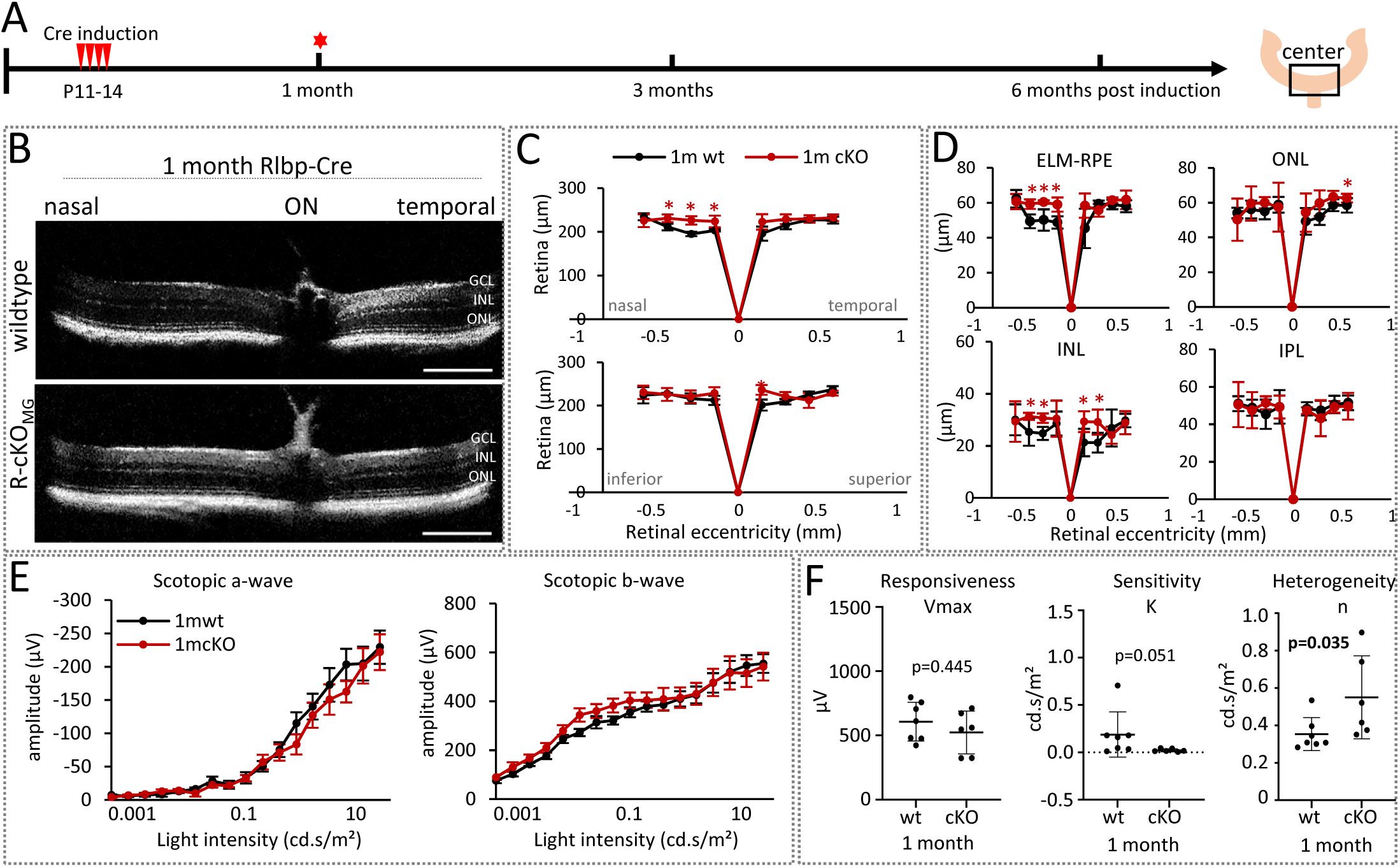
Early-stage Müller glia alterations have no impact on retinal structure and function. **A:** Experimental design. **B:** Optical coherence tomography (OCT) images of wildtype and Rlbp-Cre:Dicer cKO_MG_ (R-cKO_MG_, cKO) center retinas (1300 μm diameter) at the nasal-temporal axis 1 month after Cre induction. **C:** Spider plots of the overall retinal thickness (diameter, μm) measured at the nasal-temporal and superior-inferior axis of wildtype (n=4) and Rlbp-Cre:Dicer-cKO_MG_ mice (n=6). **D:** Spider plots of the thickness (μm) of the external limiting membrane/retinal pigment epithelium (ELM-RPE), the outer nuclear layer (ONL), the inner nuclear layer (INL), and the inner plexiform layer (IPL). **E:** Full-field scotopic electroretinogram (ERG) recordings showing a-wave and b-wave amplitudes of wildtype (n=7) and Rlbp-Cre:Dicer-cKO_MG_ mice (n=6) month after Cre induction. **F:** Estimated saturated amplitudes (Vmax, responsiveness), semi-saturation (K, sensitivity), and slope (n, heterogeneity) using the Naka-Rushton equation. OCT: mean ± S.D., ERG: mean ± S.E.M. Significant differences are indicated, Mann-Whitney-U-test: *: p ≤ 0.05. Scale bars in B: 200 µm. Layer explanation is given in Figure 1.

OCT scans of wildtype and Rlbp-Cre:Dicer-cKO_MG_ one month after Cre induction did not show any substantial abnormalities. The overall retinal thickness was about 215 μm for both conditions which equals the thickness of a normal mouse retina ^33–35^. Some nasal areas in the Rlbp-Cre:Dicer-cKO_MG_ retinas were however ∼10% thicker than the wildtype, caused by an enlarged nasal ELM-RPE area (Figure 2C-D, Supplementary Figures S1A). This suggests a possible dilatation due to either a swelling of MG endfeet (ELM enlargement), fluid build-up (or both), or a possible RPE enlargement. Furthermore, the inner nuclear layer (INL) was increased (∼10 um, (Figure 2D, Supplementary Figures S1A), an event known to occur in human retinitis pigmentosa ^36–38^.

Next, we conducted scotopic ERGs but no rod photoreceptor (a-wave amplitudes) or rod-bipolar cell (b-wave amplitudes) impairments were present one month after Dicer deletion, (Figure 2E). We then performed an in-depth analysis using the Naka-Rushton equation ^39^. This equation allows the evaluation of responsiveness of the tissue via the saturated amplitude (*V_max_*), sensitivity via the semi-saturation constant (*K*) as well as heterogeneity via slope (*n*) (Figure 2F). Although there was no difference in the responsiveness and sensitivity in the b-wave response of Rlbp-Cre:Dicer-cKO_MG_ mice, the slope was different. This indicates patchy functional impairments which are known features in early stages of retinal dystrophies with drastic photoreceptor loss such as retinitis pigmentosa ^40, 41^.

### Initial structural abnormalities, indicated by hyperreflective foci and photoreceptor layer thinning, do not result in functional impairment three months after Dicer deletion

We next analyzed the retinas of Rlbp-Cre:Dicer-cKO_MG_ mice 3 months after Dicer deletion and compared it to age-matched wildtypes (Figure 3A). No differences in the overall retinal thickness or the particular layer thicknesses were found (Figures 3B-D, Supplementary Figures S1B, S2B). However, about 40% of all mice analyzed displayed hyperreflective foci in the ELM-RPE or OPL areas (Figure 3B, arrowheads). These foci are often cell accumulations (microglia) ^19^ or small detachments caused by fluid build-up ^42–45^. Since we could not confirm microglia or macrophage accumulation in retinal cross sections, the foci could be a consequence of the ELM enlargement we found in the 1-month Rlbp-Cre:Dicer-cKO_MG_ mice. Furthermore, despite an overall preserved thickness, there was an initial reduction of the ELM/RPE region in the temporal central cKO retina (Figures 3D) indicating a possible initial photoceptor impairment. The INL in the 3-month Rlbp-Cre:Dicer-cKO_MG_ was back to normal size (Figures 3D, Supplementary Figures S1B, S2B).

**Figure 3:**
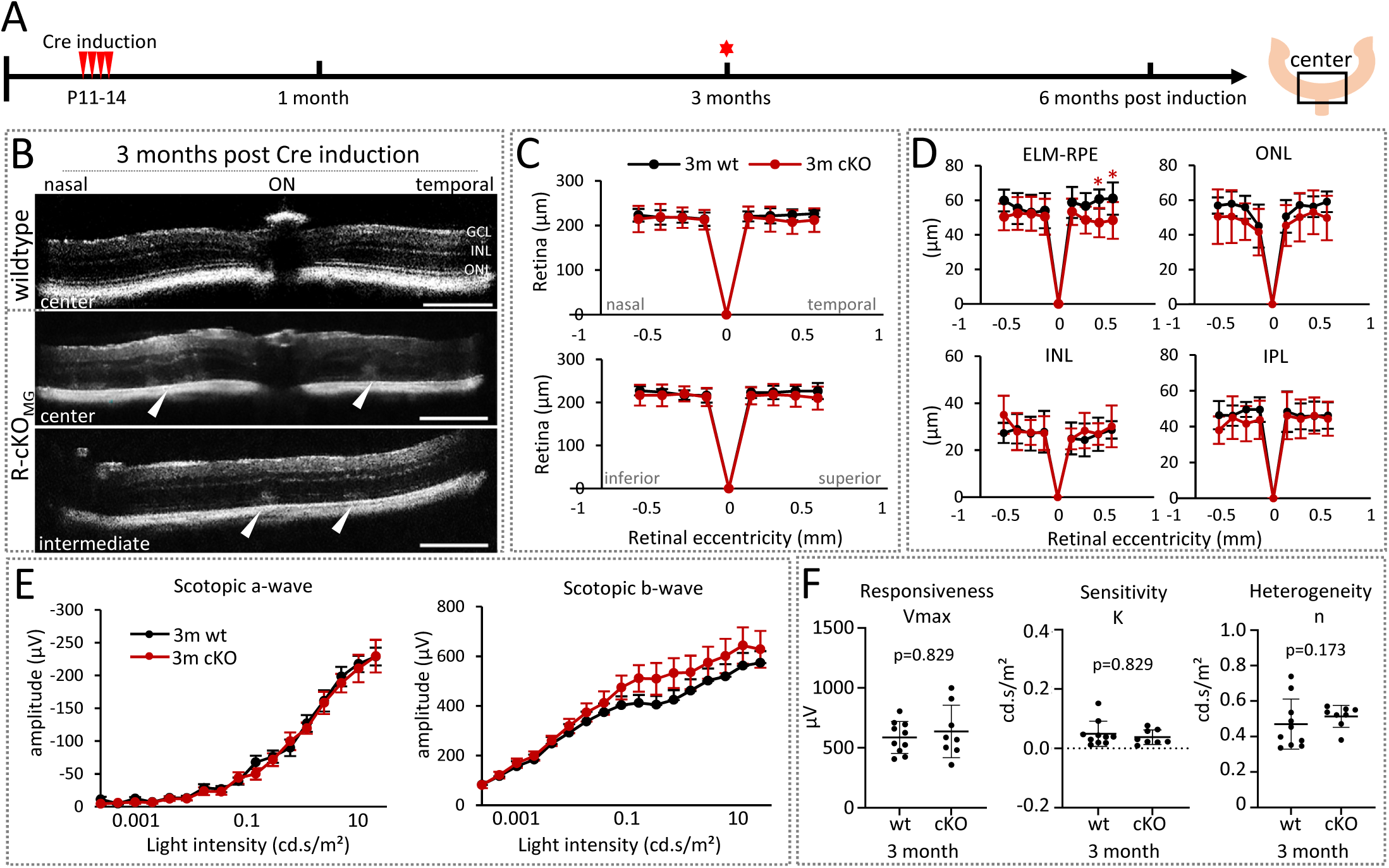
Intermediate-stage retinas display structural abnormalities but no functional impairments. **A:** Experimental design. **B:** OCT images of wildtype or Rlbp-Cre:Dicer-cKO_MG_ mice (R-cKO_MG_, cKO, center retinas (1300 μm diameter)) 3 months after Cre induction at the nasal-temporal axis. Arrowheads indicate hyperreflective foci. **C:** Spider plots of the overall retinal thickness (diameter, μm) measured at the nasal-temporal and superior-inferior axis of wildtype (n=7) and Rlbp-Cre:Dicer-cKO_MG_ mice (n=11). **D:** Spider plots of the thickness (μm) of the external limiting membrane/retinal pigment epithelium (ELM-RPE), the outer nuclear layer (ONL), the inner nuclear layer (INL), and the inner plexiform layer (IPL). **E:** Full-field scotopic electroretinogram (ERG) recordings showing a-wave and b-wave amplitudes of wildtype (n=10) and Rlbp-Cre:Dicer-cKO_MG_ mice (n=8). **F:** Estimated maximum amplitudes (Vmax, responsiveness), semi-saturation (sensitivity), and slope (heterogeneity) using the Naka-Rushton equation. OCT: mean ± S.D., ERG: mean ± S.E.M. Significant differences are indicated, Mann-Whitney-U-test: *: p≤ 0.05. Scale bars in B: 200 µm. Layer explanation is given in Figure 1.

Despite these initial structural alterations, scotopic ERGs showed no signs of neuronal impairment (Figure 3E). We however found a trend towards a slightly higher amplitude of the scotopic b-wave in the Rlbp-Cre:Dicer-cKO_MG_ retinas. Vmax, semi-saturation K, as well as the slope n were however not different from wildtype retinas. This shows not only no functional impairment at this intermediate stage but also the offset of the differences in heterogeneity found in the early stage, which appear to be transient (Figures 3F versus 2F).

### Dicer loss in Müller glia leads to significant neuronal functional loss in outer and inner retina at late stage

Next, we analyzed the late phase and conducted OCT measurements 6 months after Cre induction (Dicer/miRNA loss). We found a significantly progressed phenotype regarding pathophysiological features and functional impairment (Figures 4, Supplementary Figures S1C, S2C). The retinas of the Rlbp-Cre:Dicer-cKO_MG_ mice were thinner and displayed more hyperreflective foci. These foci seemed to be more defined (less cloudy), particularly between the ELM and RPE (Figure 4B arrows). Compared to age-matched wildtype, retinas of Rlbp-Cre:Dicer-cKO_MG_ mice displayed approximately ∼20% reduction of the overall retinal thickness (Figures 4C, D), independent of orientation. The most prominent reduction was seen in the ELM-RPE and the ONL reflecting photoreceptor loss. Interestingly, the inner layers were unchanged or slightly thicker (Figures 4D, Supplementary Figures S1C, S2C).

**Figure 4:**
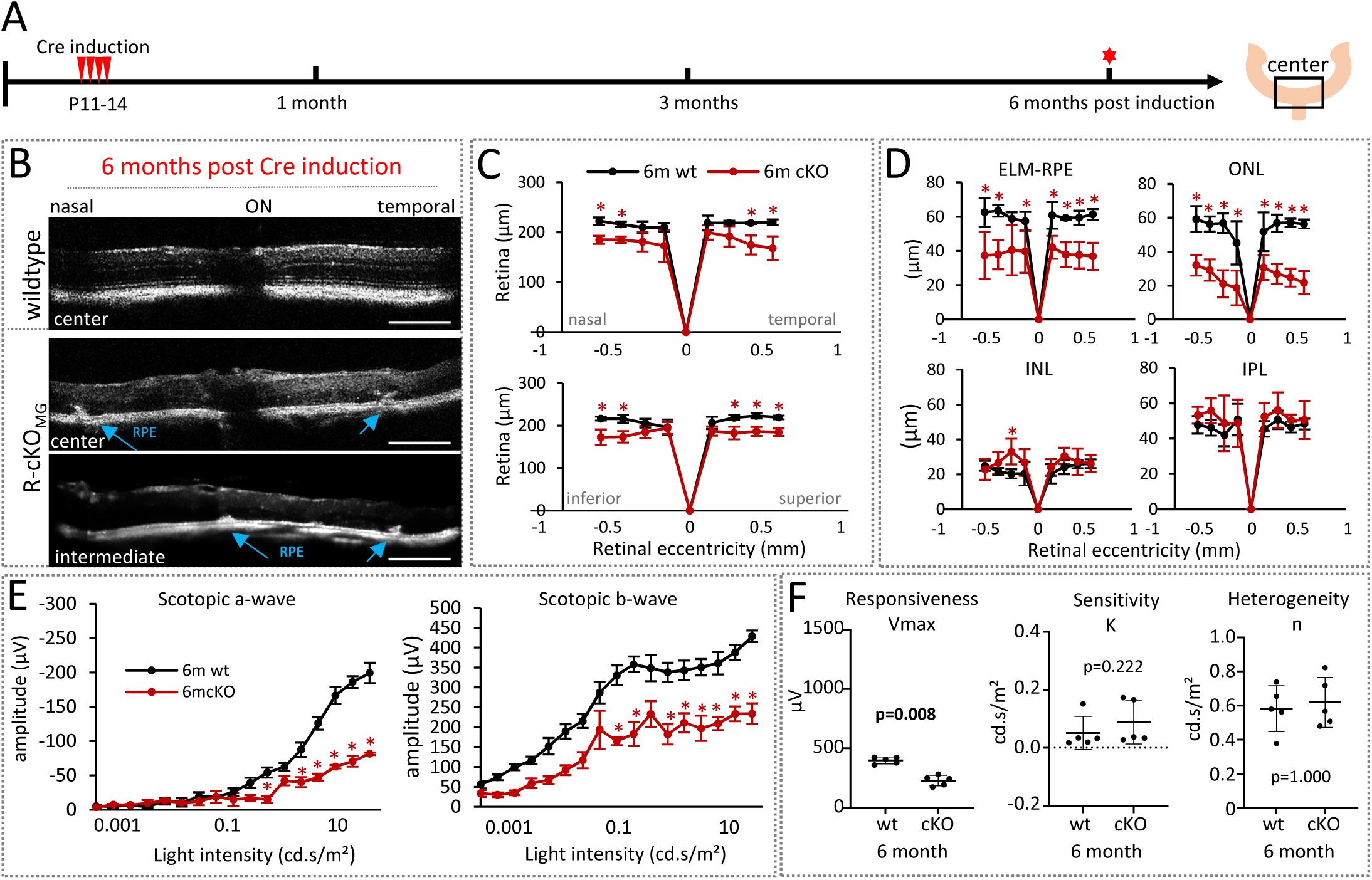
miRNA-depleted Müller glia cause photoreceptor loss and vision impairment at the late stage. **A:** Experimental design**. B**: OCT images of wildtype or Rlbp-Cre:Dicer-cKO_MG_ mice (R-cKO_MG_, cKO, center retinas (1300 μm diameter)) 6 months after Cre induction at the nasal-temporal axis. Arrows indicate hyperreflective foci. **C:** Spider plots of the overall retinal thickness (diameter, μm) measured at the nasal-temporal and superior-inferior axis of wildtype (n=4) and Rlbp-Cre:Dicer cKO_MG_ mice (n=7). **D:** Spider plots of the thickness (μm) of the external limiting membrane/retinal pigment epithelium (ELM-RPE), the outer nuclear layer (ONL), the inner nuclear layer (INL), and the inner plexiform layer (IPL). **E:** Full-field scotopic electroretinogram (ERG) recordings showing a-wave and b-wave amplitudes of wildtype (n=5) and Rlbp-Cre:Dicer cKO_MG_ mice (n=5) 6 months after Cre induction. **F:** Estimated maximum amplitudes (Vmax, responsiveness), semi-saturation (sensitivity), and slope (heterogeneity) using the Naka-Rushton equation. OCT: mean ± S.D., ERG: mean ± S.E.M. Significant differences are indicated, Mann-Whitney-U-test: *: p≤ 0.05. Scale bars in B: 200 µm. Layer explanation is given in Figure 1.

This severe pathological phenotype was accompanied by a significantly reduced scotopic a-and b-wave (Figure 4E). The Vmax (saturated amplitude) was significantly reduced, indicating reduced retinal responsiveness in the 6-month Dicer-cKO_MG_ and potential inner retinal defects (Figure 4F).

Immunofluorescent labeling using antibodies against calbindin to label horizontal cells (HCs) and amacrine cells (ACs), as well as calretinin (ACs), and acetylcholine transferase (ChAT, ACs) confirmed inner retinal defects (Figure 5). These markers also label retinal ganglion cells as well as displaced ACs in the GCL but no impairments were seen in this layer (Figure 5B-G). We found severe disruptions of the neuronal circuitry in 6-month Rlbp-Cre:Dicer-cKO_MG_ retinas with some neuronal cell bodies being located beneath the glial seal in the outer retina (Figure 5E, yellow arrow). Similar events occur during late-stage retinal remodeling in AMD or retinitis pigmentosa ^30, 46–48^. This indicates that the MG-driven retinal degeneration patterns found in the late-stage Rlbp-Cre:Dicer-cKO_MG_ mouse resemble indeed those of non-glia-based retinal degenerative models. These patterns also include the glial seal formation and the RPE alterations (Figures 5E-G, arrowheads and white arrow respectively). In particular the RPE appeared to grow inside the retina by forming quite noticeable invaginations. These invaginations resembled the shape of the hyperreflective foci seen in the OCT scans (Figure 5F, red arrows).

**Figure 5:**
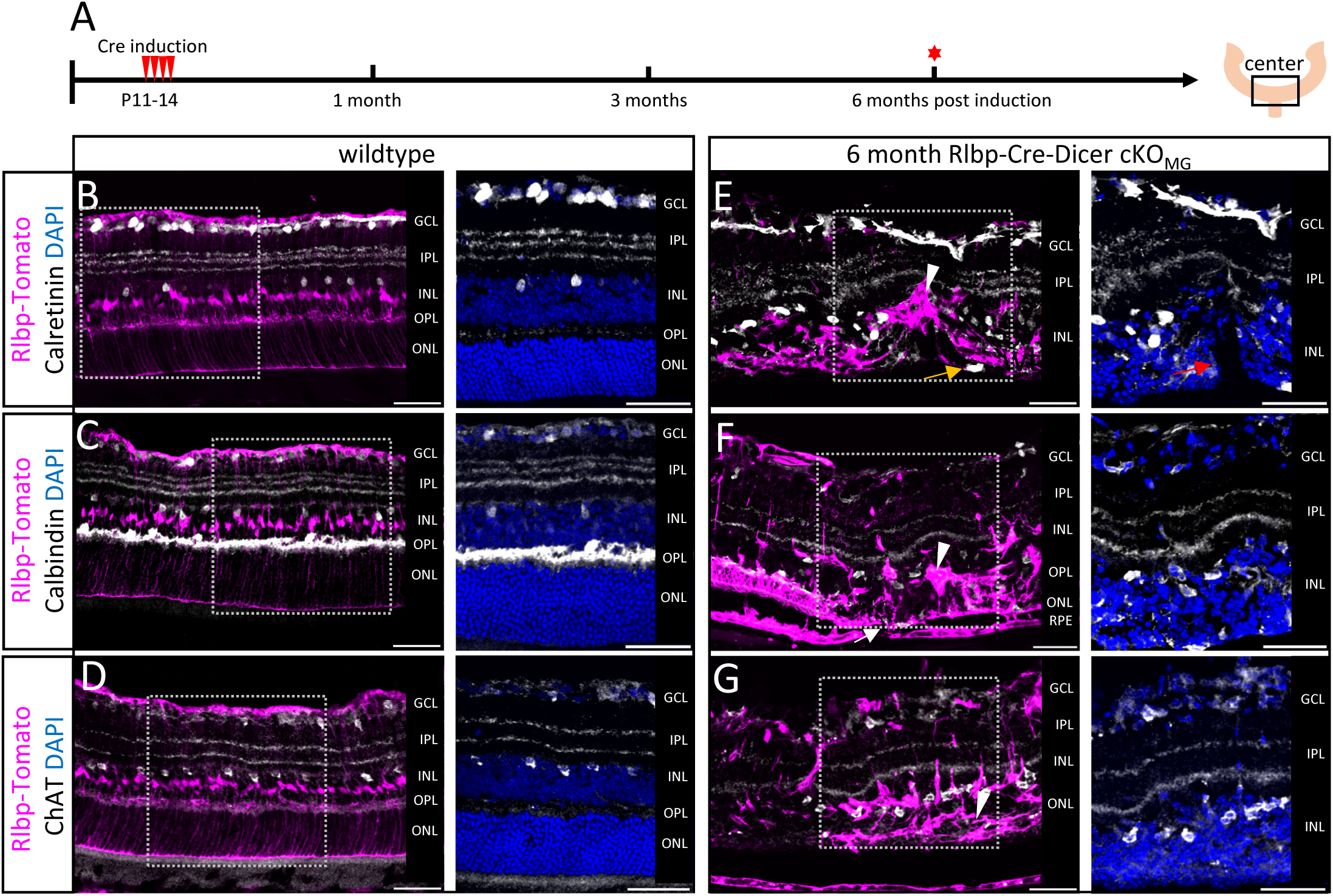
Retinal remodeling in late-stage retinas with miRNA-depleted Müller glia. **A:** Experimental design**. B-G:** Immunofluorescent labeling with antibodies against RFP (Rlbp1:Tomato), Calretinin (B,E), Calbindin (C,F) and Choline acetyltransferase (ChAT, D,G) as well as DAPI nuclear staining of center areas of wildtype (B-D) or Rlbp-Cre:Dicer cKO_MG_ mice (E-G), 6 months after Cre induction. Arrowheads in E, F, G show a glial seal, the yellow arrow in E indicates a neuronal cell body beneath the glial seal; the red arrow in E shows invagination; the white arrow in F indicates RPE-ELM interaction/ fusion. Scale bars 50 µm. Layer explanation is given in Figure 1.

### Initial rod photoreceptor degeneration is observed at 4 months post miRNA loss in Müller glia

To decipher the events happening between intermediate (3 months) and late stage (6 months) and to determine the time point of neuronal cell and neuronal functional loss we conducted OCT and ERG at 4 and 5 months after Dicer deletion (Figure 6A). OCT scans of 4-month Rlbp-Cre:Dicer-cKO_MG_ retinas showed many hyperreflective foci (cloudy appearance), predominantly in the space between the ELM and the RPE reflecting pathogenesis progression (Figure 6B, arrowheads). Thickness measurements displayed initial retinal thinning in both orientations (Figures 6C, Supplementary Figures S1D, S2D, orange graphs). The most affected layer was the ONL (photoreceptor loss) and to a lesser extent in the ELM-RPE area (unchanged compared to the 3-month time point, Figures 6D-E).

**Figure 6:**
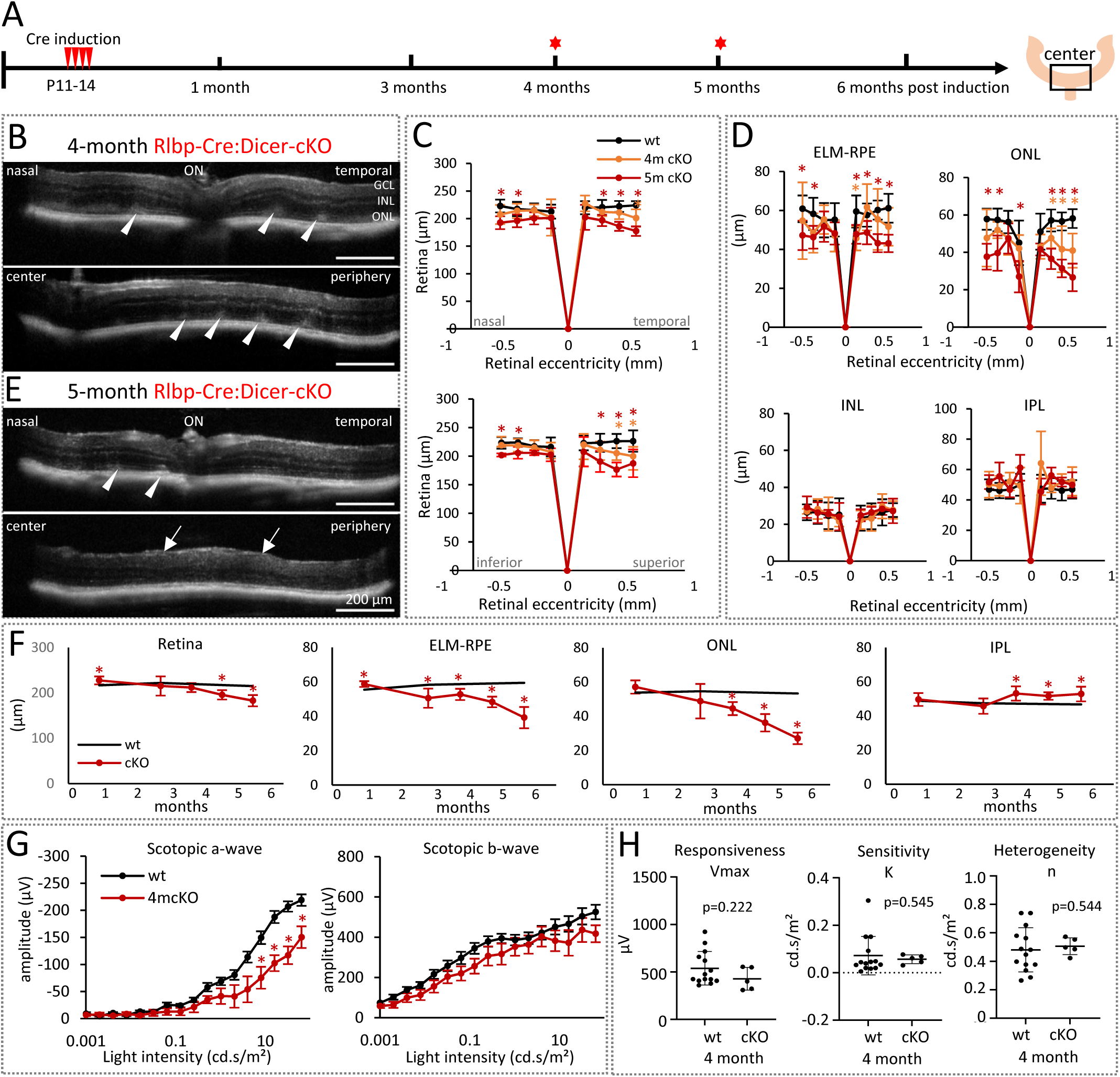
Rod functional impairments are evident 4 months after Müller glia miRNA depletion. **A:** Experimental design**. B,E:** Optical coherence tomography (OCT) images of central and peripheral retinal areas of Rlbp-Cre:Dicer-cKO_MG_ mice 4 months (B) and 5 months (E) after Cre induction. Arrowheads indicate structural impairments and arrows indicate enlarged inner nuclear layer (INL). **C:** Spider plots of the overall retinal thickness at the nasal-temporal and superior-inferior axis of wildtype (n=11), 4-month (n=5) and 5-month Rlbp-Cre:Dicer-cKO_MG_ mice (n=5). **D:** Spider plots of the thickness (μm) of the external limiting membrane/retinal pigment epithelium (ELM-RPE), the outer nuclear layer (ONL), INL, and the inner plexiform layer (IPL). **F:** Timeline plots showing the averaged thickness of Rlbp-Cre:Dicer-cKO_MG_ mice over time of the total retinal as well as specific layers. **G:** Full-field scotopic electroretinogram recordings showing a-wave and b-wave amplitudes of 4-month Rlbp-Cre:Dicer-cKO_MG_ mice (n=5) in comparison to wildtype (n=15). **H:** Estimated maximum amplitudes (Vmax, responsiveness) semi-saturation (sensitivity), and slope (heterogeneity) using the Naka-Rushton equation. OCT: mean ± S.D., ERG: mean ± S.E.M. Significant differences are indicated, Mann-Whitney-U-test: *: p ≤ 0.05. Scale bars in B,E: 200 µm. Layer explanation is given in Figure 1.

OCT scans from 5-months Rlbp-Cre:Dicer-cKO_MG_ retinas showed less hyperreflective foci in the area between the ELM and RPE (Figure 6E, arrowheads), suggesting a possible reduction of fluid build-up in this area. However, the retinal lamination appeared to be altered and in particular the IPL appeared enlarged, at least in some areas (Figure 6B arrows). The measurements of the retinal layers showed significant thinning of the total retina, especially of the ONL and the ELM-RPE area indicating progressed photoreceptor loss (Figures 6C, D, red graphs). A time course of the overall structural alterations in the Dicer-cKO (averaged measurements across the nasal-temporal and superior-inferior axis) is given in Figure 6F. The ELM-RPE area starts to decrease 3 months after Dicer deletion (∼35% decline by 6 months compared to the wildtype baseline). This ELM-RPE alteration goes hand in hand with the decrease of the ONL thickness, evident 4 months after Dicer deletion (50% reduction by 6 months compared to the wildtype baseline, Figure 6F). Conversely, the IPL increased in thickness starting 4 months after Dicer deletion and maintains this thickness until late stage despite remodeling, or perhaps because of it. The other layers remained unaffected (Supplementary Figures S3A).

ERGs conducted 4 months after Dicer deletion showed a significant reduction in the scotopic a-wave amplitude relative to wildtype, confirming a decline in functional integrity of the retina at this stage (Figure 6G). This loss of rod function correlates with the thinning in ONL and ELM-RPE observed in OCT. The scotopic b-wave, an indicator of rod-bipolar and MG function, was however not impaired, although a slight trend towards a reduced b-wave response was found (Figure 6G). The in-depth analysis of the b-wave response using Naka-Rushton equation showed that responsiveness (Vmax), sensitivity (K), and heterogeneity (slope, n) were not altered at this stage (Figure 6H). The time course of the scotopic a- and b-wave ranging from 1 - 6 months after Dicer deletion and the time course of the Naka-Rushton parameters are given in Supplementary Figures S3B-C showing the first functional rod deficit 4 months after Dicer deletion.

### The loss of cone function precedes the loss of rod function

Since MG play an essential role in cone pigment recycling, we next measured the photopic b-wave response 3, 4, and 6 months after Dicer deletion to analyze cone health in the cKO mouse (Figure 7A). A photopic a-wave is not detectable in rodents due to a small cone population^49, 50^. We found that cone loss precedes rod loss: 3-months after Dicer deletion the photopic b-wave amplitude in the cKO was reduced by ∼20% compared to age-matched wildtypes and further dropped with age (Figure 7B-D). Immunofluorescent labeling with antibodies against M-Opsin showed less dense cone outer segments in cKO retinas that also appeared scattered/fragmented, validating the functional deficit (Figure 7E-H,H’, arrowheads).

**Figure 7:**
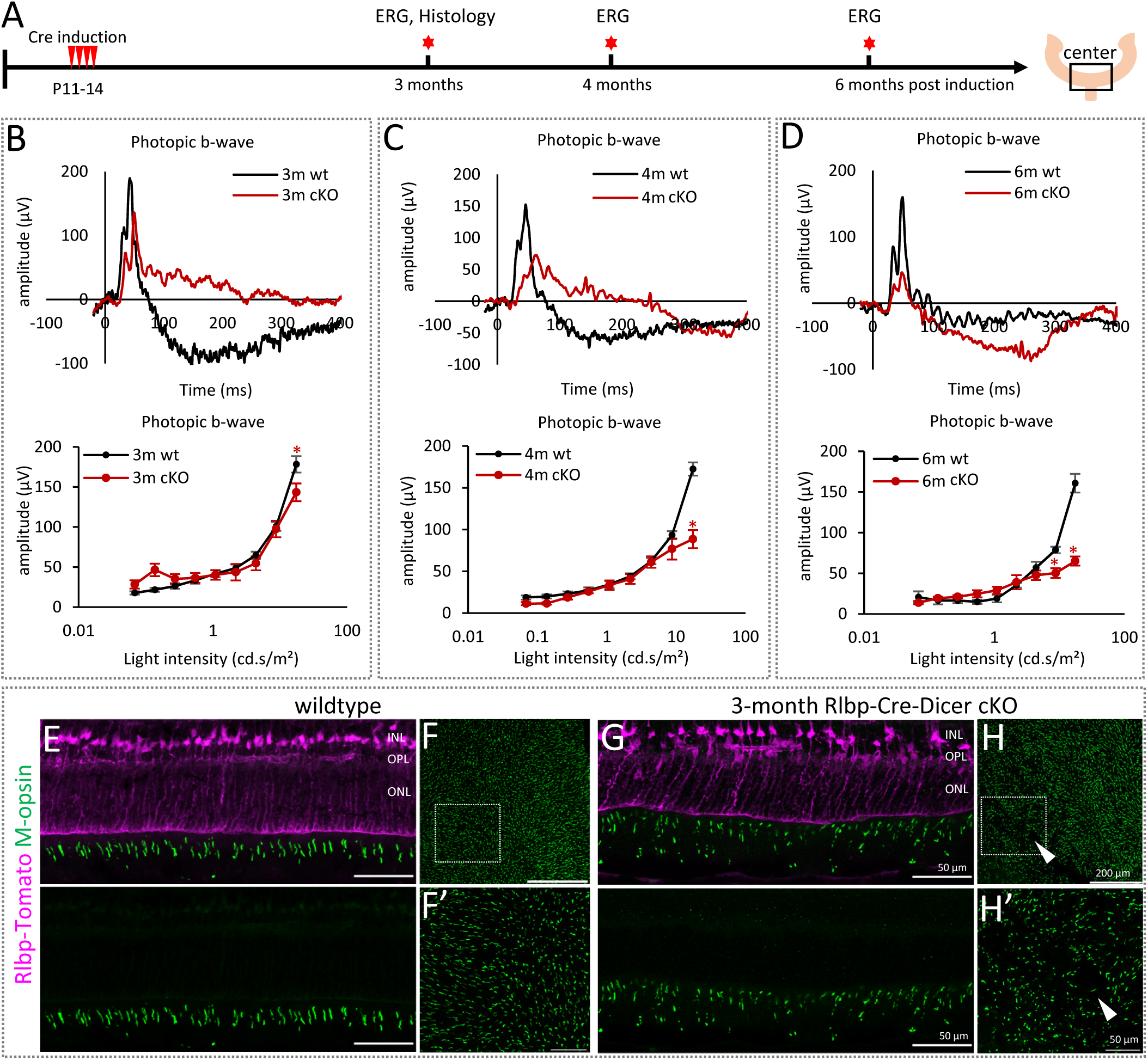
Rlbp-Cre:Dicer-cKO mice display loss of cone function 3 month after Cre induction. **A:** Experimental design. **B-D:** Full-field photopic electroretinogram (ERG) recordings showing representative ERG waveforms, b-wave intensity-amplitude plots of wildtype and Rlbp-Cre:Dicer cKO_MG_ mice 3-month (B: wt: n=10 vs. cKO: n=8), 4-month (C: wt: n=15 vs. cKO: n=5) and 6-month (D: wt: n=5 vs. cKO: n=5). **E-H:** Immunofluorescent labeling of cross-sections (E, G) and flat mounts (F/F’,H/H’) with antibodies against RFP (Rlbp1:Tomato) and M-opsin (medium wavelength cone opsin) of center areas of wildtype (E-F) and 3-month Rlbp-Cre:Dicer cKO_MG_ mice (G-H). Insets in F and H are shown in F’ and H’. Arrowheads indicate areas with reduced and scattered M-opsin expression. Mean ± S.E.M., significant differences are indicated, Mann-Whitney-U-test: *: p ≤ 0.05. Scale bars in E,G,F’,H’: 50 µm, in F, H: 200 µm. Layer explanation is given in Figure 1.

### The loss of miRNAs in the RPE causes partial RPE alteration

Since Rlbp1 is not only expressed in MG, but also in the RPE, a contribution of a malfunctioning RPE to the observed rod loss in Dicer-cKO_MG_ mouse cannot be ruled out. We therefore labeled flat-mounted RPE with zonula occludes 1 (Zo1), a marker for tight junctions of the RPE ^51^ and Otx2, a marker known to be expressed in mature RPE ^52, 53^. Wildtype and cKO RPE displayed a mosaic expression, in accordance to previous reports ^22, 54^. Some animals had about 70% reporter induction, while others only had about 40%. Wildtype RPE cells as well as reporter negative cells in the cKO were healthy, displaying defined Zo1 labeled membranes and one or two Otx2+ nuclei in the center of the cell (Figures 8A-A’).

**Figure 8:**
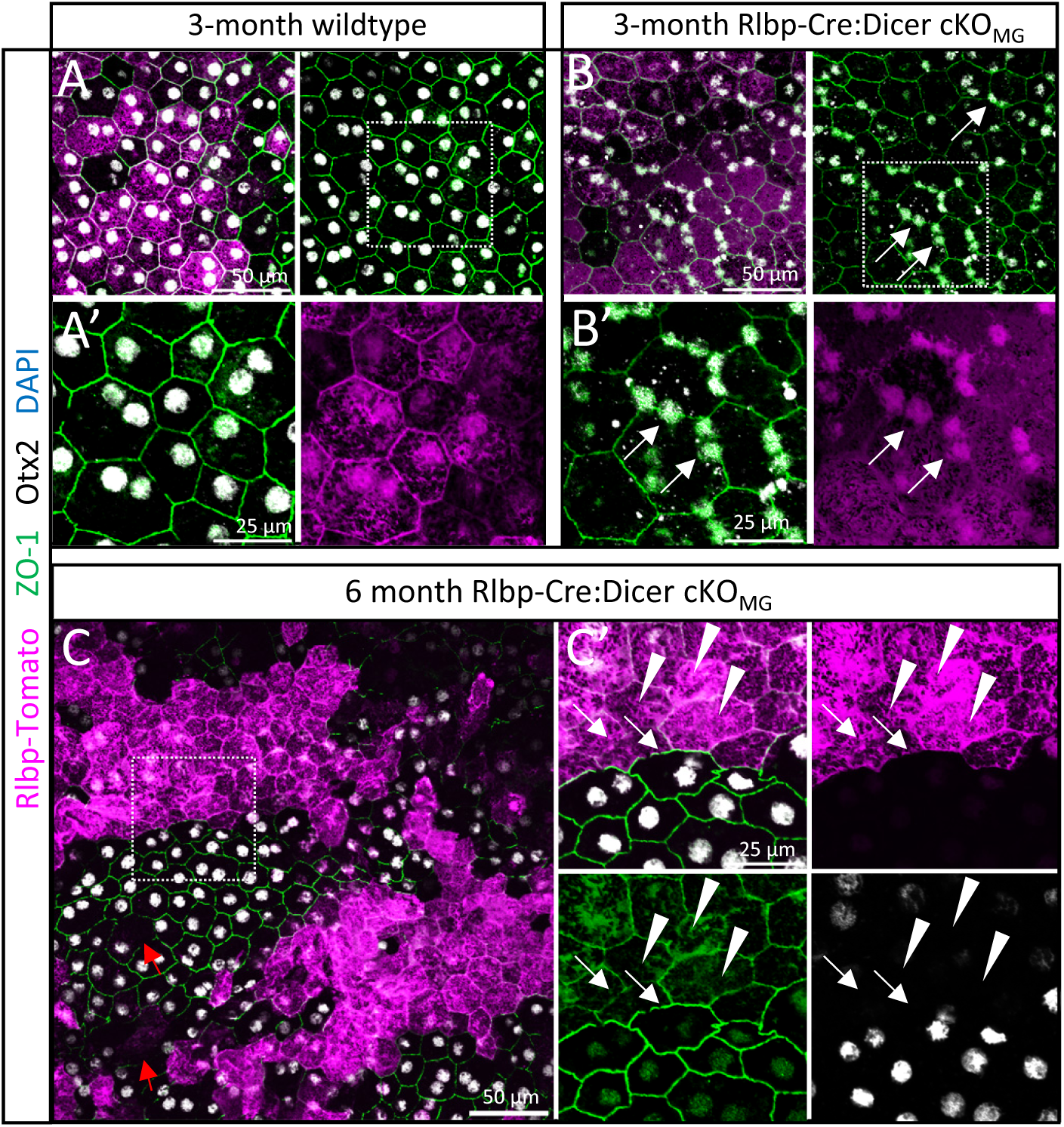
RlbpCre:Dicer-cKO mice have partially impaired RPE. Immunofluorescent labeling with antibodies against RFP (Rlbp1:Tomato) ZO-1, Otx2 as well as DAPI nuclear staining RPE flat mounts of 3-month wildtype (A), 3-month RlbpCre:Dicer-cKO (B) and 6-month RlbpCre:Dicer-cKO (C). Insets in A-C are shown in A’-C’. Arrows in B/B’ show altered nuclei location in the 3-month RlbpCre:Dicer-cKO. Arrows in C’ show enlarged RPE cells or RPE cells that lost their ZO-1+ membrane. Arrowheads in C’ show RlbpCre:Dicer-cKO RPE cells with reduced ZO-1 and Otx2 expression. Scale bars in A-C: 50 μm; in A’-C’: 25 μm.

Three-month cKO RPE cells however, had less intense Zo1 and Otx2 labeling and dislocated nuclei (towards the cell membrane, Figures 8B-B’ arrows). These dislocated nuclei appeared to be smaller and were mostly arranged in pairs. Some cells seem to have several nuclei, suggesting either cell division or a potential nuclear migration. Six-month-cKO RPE cells displayed a more progressed phenotype: many cells displayed very weak Zo1 expression in the cell membrane and faint Otx2+ or Otx2-nuclei (Figures 8C-C’ arrowheads) indicating impaired RPE. Interestingly, the areas in the cKO RPE in which tdTomato (reporter)-negative “normal” RPE cells were found. They displayed mostly wildtype-like Zo1 and Otx2 expression pattern. However, some of these normal (reporter-negative) RPE cells appeared to be enlarged with partly absent Zo1 membranes (Figure 8C arrows). This may suggest a possible negative influence of adjacent cKO RPE cells. Overall, the partial alteration of the RPE in the Rlbp-Dicer-cKO mouse very likely contributed to observed retinal (rod) degeneration.

### Glast-Cre:Dicer-cKO mice display a similar disease phenotype but with more persevered retinal structure

To assess the extent of RPE contribution to the observed phenotype in the Rlbp-Cre:Dicer-cKO mouse, we created the Glast-Cre:Dicer-cKO_MG_ mouse with no reporter expression in the RPE^54^. Glast, i.e. glutamate aspartate transporter (encoded by Slc1a3 and also known as EEAT1) and is expressed in brain astrocytes ^55^ and MG. We used the same experimental paradigm and validated the reporter induction in 3-month old mice (Supplementary Figures S4). RPE and retinal astrocytes were reporter negative in accordance to previous reports ^29, 54^. Retinal cross sections and flat mounts showed similar induction patterns in MG compared to Rlbp-Cre. The efficiency was however quite heterogenous and a little less efficient (Glast-Cre: 70-90% vs. Rlbp-Cre: ∼98% reporter+ MG of total MG).

Next, we conducted OCT and ERG analysis of the Glast-CreERT wildtype and cKO mice 3 and 6 months after Cre induction (Dicer deletion) to analyze structure and function (Figure 9A). OCT scans showed that Glast-Cre wildtype mice were not different form Rlbp-Cre or any other wildtype strain. In the 3-month Glast-Cre:Dicer-cKO_MG_, the overall retinal architecture was normal but hyperreflective foci were present in the ELM-RPE region (Figure 8B, arrowheads, Supplementary Figures S5A-C). That suggests that these hyperreflective foci are primarily caused by MG dysfunction and not by RPE impairment. The retinas of 6-month Glast-Cre:Dicer-cKO_MG_ mice were quite heterogeneous (probably due to less efficient induction), had thinner retinas than the wildtypes, and displayed more hyperreflective foci between the ELM and RPE compared to the Rlbp-Cre:Dicer-cKO_MG_ retinas at this stage (Figure 9D-E, arrowheads). This thinning was due to a reduction of the ELM-RPE area and the ONL, hence affecting photoreceptors. Overall, although the Rlbp-Dicer-cKO strain appeared more affected (ONL), both cKO-strains resulted in similar thinning (Figure 9E-F, Supplementary Figures S5D, E).

**Figure 9:**
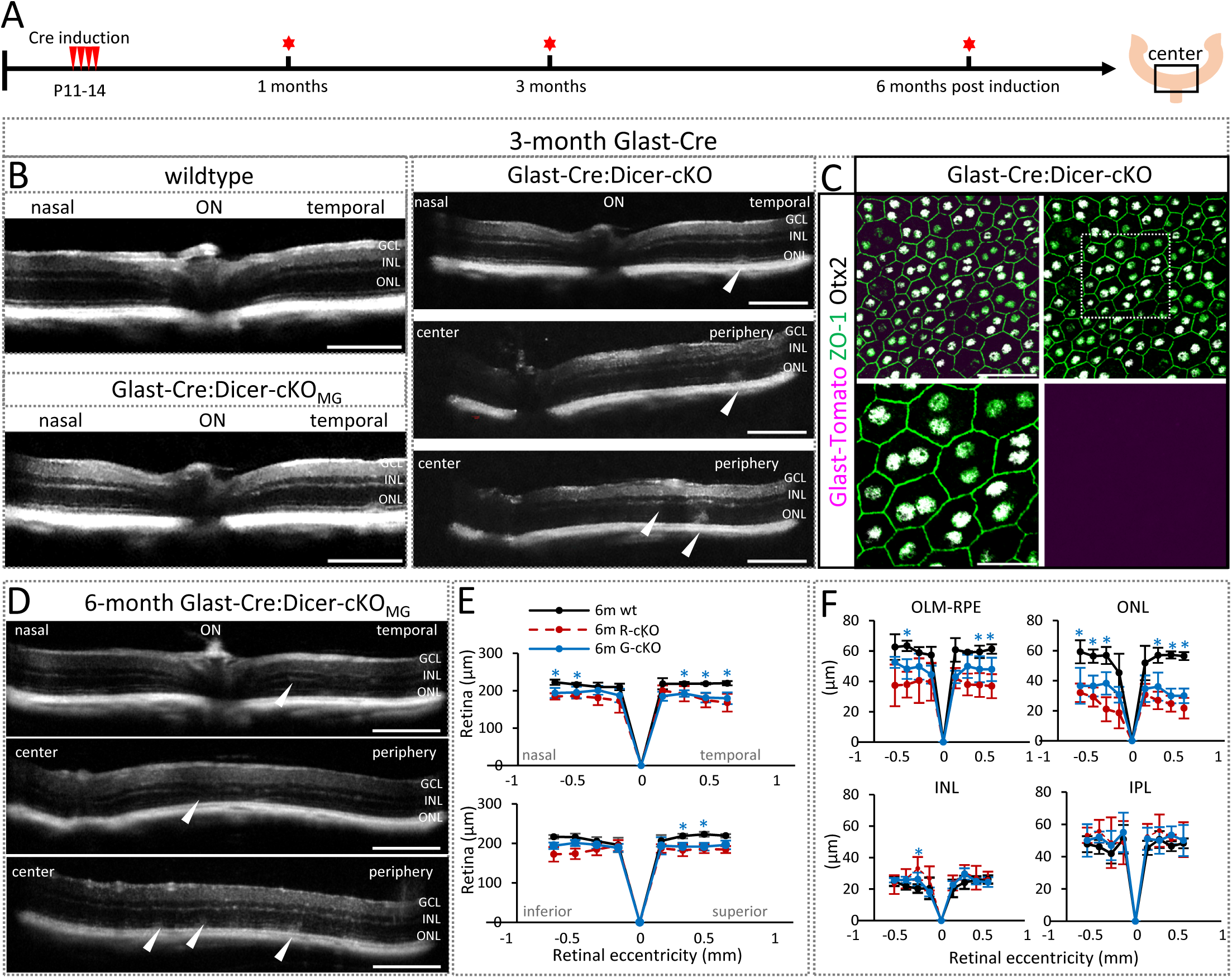
Glast-Cre:Dicer-cKO mice display hyperreflective foci and retinal thinning. **A:** Experimental design**. B:** Optical coherence tomography (OCT) images of wildtype and or Glast-Cre:Dicer-cKO_MG_ center at the nasal-temporal axis 3 months after Cre induction. **C:** Immunofluorescent labeling of Glast-Cre:Dicer-cKO RPE flat mounts 3 months after Cre induction with antibodies against RFP (Glast:Tomato), ZO-1, Otx2 as well as DAPI. **D**: OCT images of Glast-Cre:Dicer-cKO_MG_ center at the nasal-temporal axis 6 months after Cre induction. Arrowheads in B and D indicate structural impairments. **E:** Spider plots of the overall retinal thickness at the nasal-temporal and superior-inferior axis of wildtype (n=5) and Glast-Cre:Dicer-cKO_MG_ mice (n=6) 6 months after Cre induction, in comparison to age-matched Rlbp-Cre:Dicer cKO_MG_ mice (n=7). **F:** Spider plots of the thickness (μm) of the retinal layers for wildtype and Glast-Cre:Dicer-cKO_MG_ mice, in comparison to Rlbp-Cre:Dicer-cKO_MG_ mice. Mean ± S.D. Significant differences are indicated, Mann-Whitney-U-test: (*) wt-cKO comparison, (+) cKO-cKO comparison: p≤ 0.05. Scale bars in B-D: 200 µm; in inset: 25 µm. Layer explanation is given in Figure 1.

### Glast-Cre:Dicer-cKO display impaired cone function but more persevered rod function

Since the first neuronal dysfunctions in the Rlbp-Cre:Dicer-cKO_MG,_ mouse concerned cones, we next conducted photopic ERG recordings in the Glast-Cre:Dicer-cKO_MG,_ mouse 3 and 6 months after Dicer deletion. Despite unaltered retinal structure, we found cone impairment 3 months after Dicer deletion, further progressing over time (Figures 10A,B). M-Opsin expression appeared scattered and fragmented displaying the same phenotype as the Rlbp-Cre:Dicer-cKO_MG,_ mouse (Figures 10C,D vs. 7). Since both MG Dicer-cKO_MG,_ strains resulted in the same outcome, cone functional loss appears to be due to molecular (miRNA) alterations in MG.

**Figure 10:**
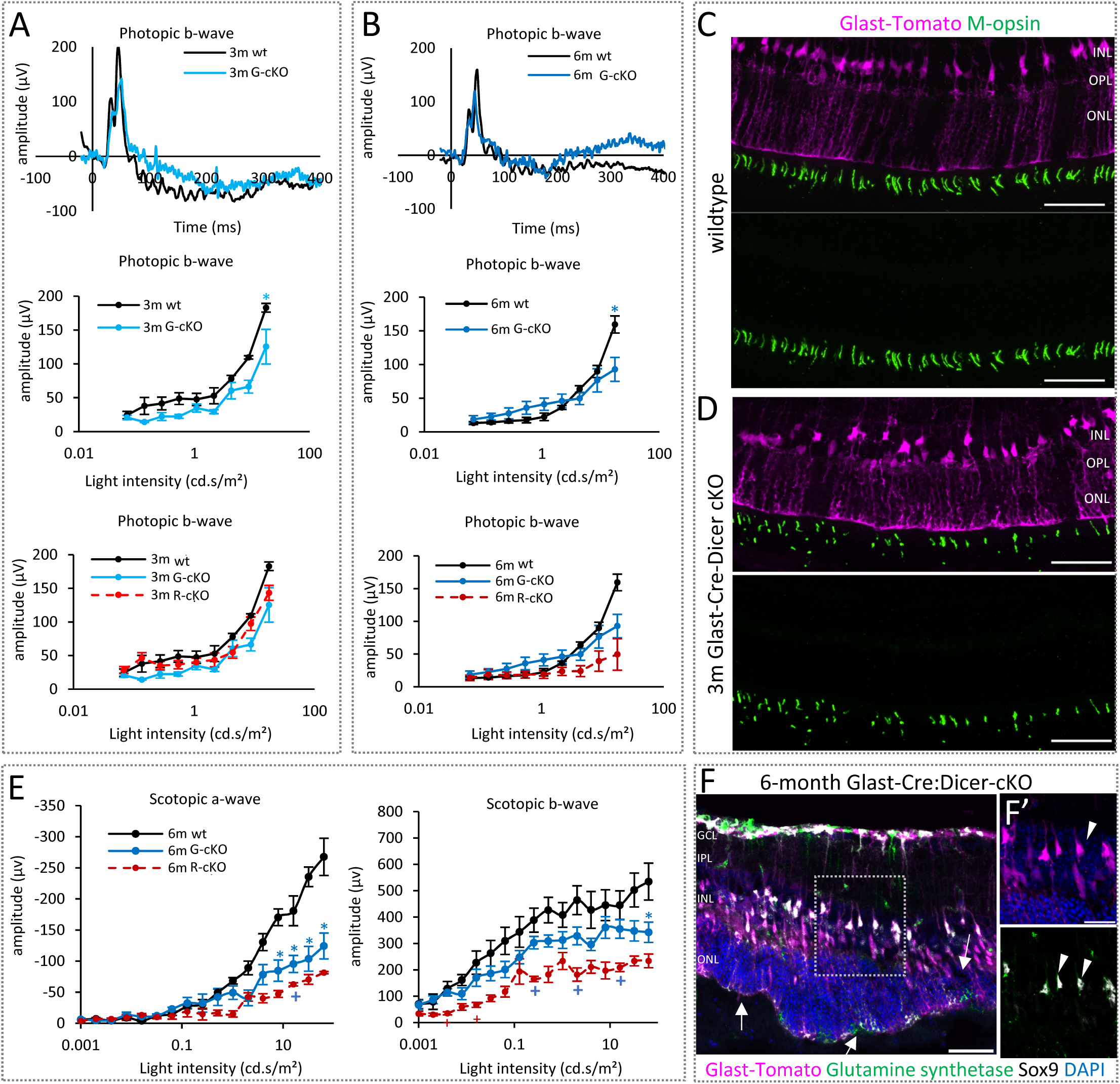
Glast-Cre:Dicer-cKO mice show loss of cone function 3 months after Dicer deletion. **A-B**: Full-field photopic electroretinogram (ERG) recordings showing representative ERG waveforms, b-wave intensity-amplitude plots of 3-month Glast-Cre:Dicer-cKO_MG_ mice (n=5) in comparison to wildtype (n=5, A) and 6-month Glast-Cre:Dicer-cKO_MG_ mice (n=6) in comparison to wildtype (n=5, B). **C-D**: Immunofluorescent labeling of retinal cross-sections with antibodies against RFP (Glast:tdTomato) and M-opsin (medium wavelength cone opsin) of center areas of wildtype (C) and 3-month Glast-Cre:Dicer-cKO_MG_ mice (D). **E:** Full-field scotopic electroretinogram recordings showing a-wave and b-wave amplitudes of Glast-Cre:Dicer-cKO_MG_ (n=6) in comparison to wildtype (n=5) and Rlbp-Cre:Dicer-cKO_MG_ (n=5). **F-F’:** Immunofluorescent labeling of central retinal areas of the Glast-Cre:Dicer-cKO_MG_ mouse 6 months after Dicer deletion, with antibodies against RFP (Glast:Tomato), Glutamine Synthetase (GS) and Sox9, as well as DAPI nuclear staining. Arrows in F show a disturbed ELM, arrowheads in F’ display GS reduced MG. Scale bars in C,D and F: 50 µm, in F’: 25 µm. Mean ± S.E.M., significant differences are indicated, Mann-Whitney-U-test: (*) wt-cKO comparison, (+) cKO-cKO comparison: p≤ 0.05. Layer explanation is given in Figure 1.

Scotopic ERG recordings 3 and 4 months after Dicer deletion showed no rod photoreceptors impairment, contrary to the observations in the Rlbp-Cre:Dicer-cKO_MG,_ mouse (Supplementary Figures S6 A-B). Naka-Rushton analysis showed also no alterations except an increased Vmax (responsiveness) in the 4-month Glast-Cre:Dicer-cKO_MG,_ mouse compared to the wildtype. As mentioned before, a similar trend was seen in the Rlbp-Cre:Dicer-cKO_MG_ mouse (Supplementary Figure S6 B vs. 3E).

6-month Glast-Cre:Dicer-cKO_MG,_ retinas displayed rod dysfunction (scotopic a-wave) leading also to initial functional loss in the inner retina (scotopic b-wave, Figure 10E, Supplementary Figures S6 C). Overall, the scotopic functional loss in the 6-month Glast-Cre:Dicer-cKO_MG_ mouse resembled the functional loss found in the 6-month Rlbp-Cre:Dicer-cKO_MG_ mouse. However, the inner retina (rod bipolar cells) in the 6-month Glast-Cre:Dicer-cKO_MG_ were less affected (unchanged Vmax, Supplementary Figure S6 C). Histological analysis showed that Glast-Cre:Dicer-cKO_MG_ retinas too had areas with massive photoreceptor loss and ELM impairments (Figure 10F-H, arrows). In addition, also MG in the 6-month Glast-Dicer-cKO had reduced or absent glutamine synthetase labeling (Figure 10F’, arrowheads).

## Discussion

Many retinal dystrophies are not fully understood yet. While retinitis pigmentosa or macular degenerations have predominantly genetic causes primarily affecting the RPE and photoreceptors, there is growing evidence that MG are not only responders to tissue changes, but also contribute to diseas progression ^56–60^ ^61^ or might even participate in the initiation of degenerative events ^19, 20, 62–64^.

MG have a specific set of miRNAs different from that of neurons ^29^. The loss of these MG-specific miRNAs in young glia results in abnormal glial behavior and a slow but severe photoceptor loss with subsequent retinal remodeling ^9^, as it occurs in retinitis pigmentosa or AMD ^46, 65–68^. Therefore, any pathologies seen after manipulation are caused by malfunctioning MG, making this mouse a very interesting model for studying MG-induced retinal disease phenotypes. The onset and the order of events of the observed neurodegeneration as well as whether the RPE contributes to the phenotype was not known and investigated in this study.

Here we show that MG-dysfunction, caused by the deletion of Dicer and mature miRNAs in the glia, leads first to impairments of cone health (summarized in Figure 11). First pathological indications are found in area comprising the ELM-RPE. Structural and functional cone impairment were evident in two different MG-reporter lines, i.e. Rlbp-Cre or Glast-Cre, 3 months after manipulation, validating this robust phenotype. Retinas of both strains displayed hyperreflective foci, which are considered indicators for disease ^69^, in particular in AMD attributed to ectopic RPE ^70–72^. Our Dicer-cKO model now suggests that hyperreflective foci can also be attributed to MG malfunction. Furthermore, cone impairment was followed by rod impairment, which was more accelerated in the Rlbp-Cre-driven strain, due to partial RPE malfunction. Therefore, the Glast-Cre allows the study of MG-driven events, while Rlbp-Cre strain allows additionally the study of RPE-related events and RPE-MG interactions.

**Figure 11:**
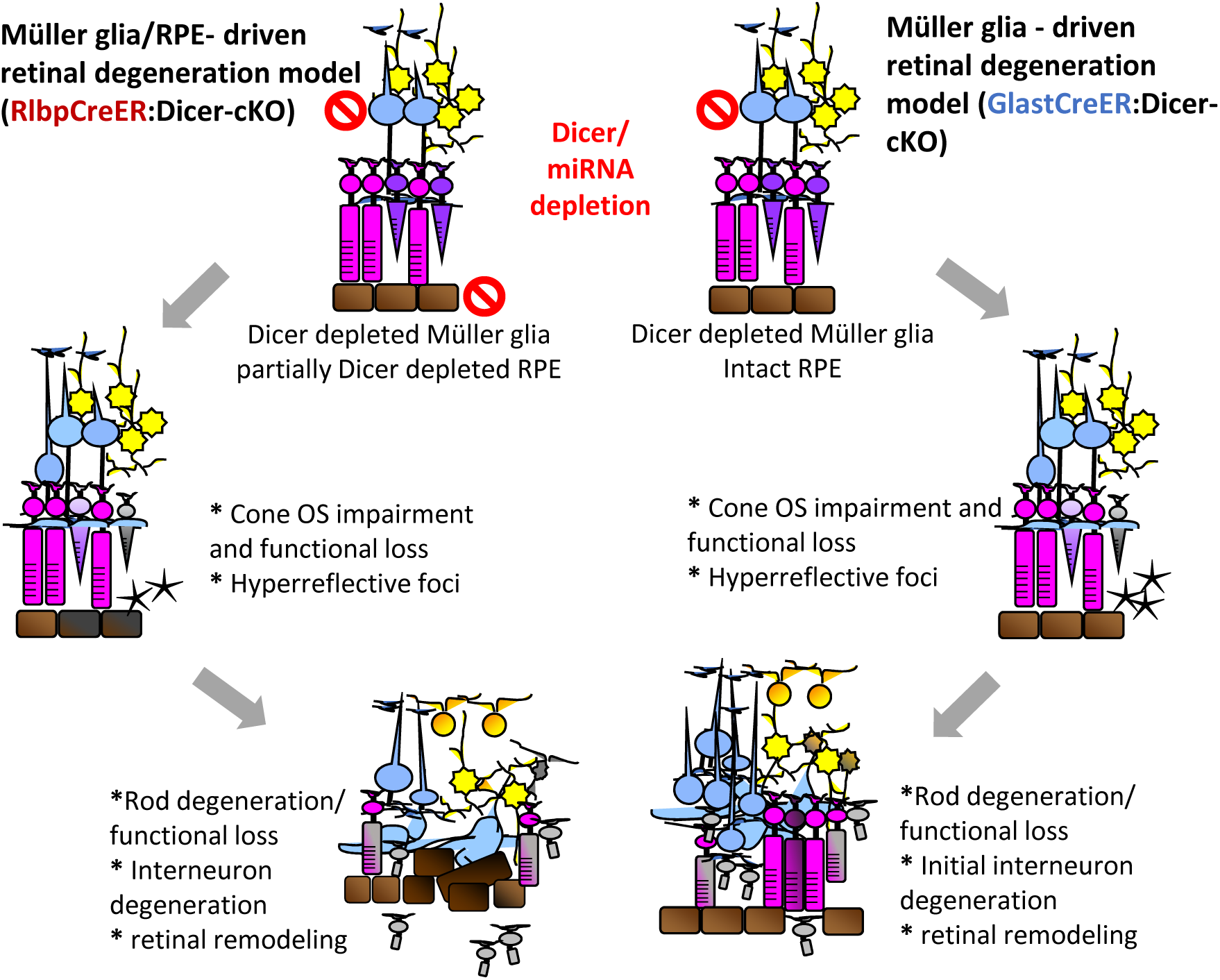
Dicer-depleted Müller glia lead to cone functional loss with subsequent rod loss. Schematic overview of the major events happening in the MG/RPE-driven retinal degeneration model (RlbpCreER:Dicer-cKO) and the MG-driven retinal degeneration model (GlastCreER:Dicer-cKO) emphasizing the shared pathological features: cone outer segment impairment and cone functional loss and hyperreflective foci in the external limiting membrane-retinal pigment epithelium (ELM-RPE) at the intermediate state and rod degeneration/ rod functional loss and interneuron degeneration leading to retinal remodeling at late stage. Additional features evident in both models include ELM breakage and glial seal formation. The RlbpCreER:Dicer-cKO model displayed furthermore impaired RPE contributing to rod loss and accelerating the overall pathological phenotype.

### First pathological indications after conditional loss of Dicer and mature miRNAs in Müller glia are found in the ELM-RPE

As early as one-month after Dicer deletion, we found a thicker ELM-RPE. The ELM, formed by tight and adherens junctions of MG and photoreceptors plays a substantial role in disease manifestation and progression. An intact ELM is a requirement for an intact ellipsoid zone and the integrity of both structures is vital for the preservation of normal vision. The ELM represents a barrier for macromolecules and is a structure formed by junctional complexes between inner photoreceptor segments and the outer processes of MG ^73–76^. Alterations in the ELM with subsequent changes in the ellipsoid zone have been reported in diabetic retinopathy ^77–81^ and Stargardt disease ^61, 82, 83^. In the form of diabetic retinopathy characterized by diabetic macular edema, it was reported that MG undergo hypertrophy. Their swollen endfeet lose their connection to the photoreceptors inner segments due to a decline in occludin ^76^, a transmembrane protein that contributes to tight junctions stabilization and optimal barrier function ^84^. This disruption then leads to the accumulation of intraretinal fluid, primarily in the inner and outer plexiform layers and manifests as retinal thickening and hyperreflective loci.

Furthermore, it is suggested that VEGF alters tight junctions and promotes vascular permeability in the central nervous system including the neural retina. Although the mechanisms of this barrier regulation are not yet fully understood, it was reported that VEGF induces phosphorylation-dependent occludin ubiquitination which is necessary for increased permeability to macromolecules and ions ^85^. MG are known to express VEGF ^86, 87^ and therefore have been suspected to contribute to the manifestation of the events seen in diabetic retinopathy ^88–90^. The suspicion that MG are one or even the driving component in these diseases was confirmed by MG ablation studies. The partial elimination (absence) of MG resulted in the same pathophysiological events ^19, 20^. In our study, which is solely based on Dicer deletion with subsequent mature miRNAs depletion in the glia, we found these hallmarks suggesting that MG miRNAs might regulate these events. It was shown that miRNAs that directly target VEGF (3’UTR) include miR-16, miR-20a, miR-29b, miR-125, miR15c, and miR-200c ^91–96^ and some of them are expressed in MG ^29^.

Moreover, MG have been shown to actively contribute to absorbing excess fluid from the subretinal space via the water pore aquaporin 4 (Aqp4) ^97, 98^. Two miRNAs, miR-29b and miR-320a were identified as aquaporin4 suppressors. miR-29 was found downregulated after stroke/cerebral ischemia and its overexpression reduced the disruption of the blood brain barrier after ischemic stroke by downregulating aqp4 ^99^. Similarly, overexpression of miR-320a in the retina resulted in aquaporin 4 downregulation and attenuated hypoxia-induced injury including retinal edema ^100^. A compromised MG function due to miRNA depletion might also cause an alteration in aquaporin expression causing liquid accumulation.

As mentioned before, ELM disruption can lead to fluid accumulation resulting in retinal thickening and hyperreflective loci. We found in our two independent MG Dicer cKO models (Rlbp-Cre and GlastCre), hyperreflective foci in the ELM/RPE area 3 months after Dicer/miRNA depletion. Hyperreflective foci belong to pathological features found in diabetic retinopathy/diabetic macular edema ^42^, retinitis pigmentosa ^43–45^, Stargardt disease ^61, 82, 83^ and macular telangiectasia ^101^ and are a consequence of the ELM impairment. Hyperreflective foci can also be detected in age-related macular degeneration (AMD) where they represent deposits rather than edema ^70, 102, 103^. Since these foci were found in both of our cKO models (Rlbp-Cre and Glast-Cre) it stands to reason that they are predominantly a consequence of MG malfunction and less due to RPE malfunction. This conclusion is in accordance with two previous independent MG ablation studies in which the RPE was not affected ^19, 20^. This suggests that some retinal dystrophies and degenerative diseases, in particular the ones with unknown (non-genetic) origin, might be caused by imbalanced MG-miRNAs. Furthermore, the fact that the same pathophysiological ELM-ellipsoid zone alterations were seen in animal models of retinal dystrophies ^45, 104^ and ischemia ^105^ which are models in in which solely neurons are targeted, further supports the hypothesis that the driving force for these events are the glia.

### A decline in cone photoreceptor function precedes rod dysfunction

Muller glia have a plethora of functions in the healthy retina, one of them is cone pigment recycling ^18, 106–109^. We found impaired cone photoreceptor structure and function in both MG-specific Dicer-cKO strains, suggesting a central role for MG miRNAs in maintaining cone health in the retina. Cones are the photoreceptors involved in daylight and color vision and maintaining cone health is crucial to maintaining proper visual function. Loss of cones and cone function is a feature of some retinal diseases such as AMD ^110, 111^, Leber’s Congenital Amaurosis^112^ ^116^, and Stargardt disease ^61, 113, 114^. The cell type predominantly driving the retinal visual cycle are the MG ^107, 115^. MG are generating 11-cis retinol from the all-trans form via the enzyme cellular retinaldehyde binding protein (CRALBP), encoded by Rlbp1 ^116–119^. Hence MG play a substantial role in visual pigment regeneration. Although the direct link between MG miRNAs and cone function in this model remains unknown and requires a subsequent investigation, it has been shown that loss or reduction of Cralbp expression MG results in dislocated M-opsin and reduced cone function ^109, 120^. Furthermore, it has been shown that peripheral regions in the RD1 retinitis pigmentosa mouse model harbor functional cones and that their prolonged survival is ensured by retinoic acid signaling of peripheral MG ^121^. This suggests that MG-miRNAs might play a role in regulating the MG-driven visual cycle.

### Dicer/miRNA loss in MG leads to rod loss with partial Dicer deletion in the RPE accelerating the process

Both MG-specific Dicer cKO strains resulted in rod loss accompanied by functional impairments that later on affected the inner retina. The overall progression of rod loss was very slow but the pattern resembled the events found in rodent models of retinitis pigmentosa ^122–124^ and in the early stages of human forms of retinitis pigmentosa ^125^ caused by genetic defects. This not only suggests that MG contribute to the retinitis pigmentosa phenotype but also that MG miRNAs seem to regulate their behavior. Furthermore, we here utilized the Naka-Rushton equation, an analysis predominantly performed on patient ERGs, that allows an in-depth analysis of the b-wave with regard to responsiveness (Vmax), sensitivity (K) and heterogeneity (slope) ^39–41^. Overall our Dicer-cKOs displayed similarities comparable to observations made in patient ERGs ^125^. These similarities include a change in slope at early stages indicative for a patchy onset of functional loss ^40, 41^. This is followed by a reduction in responsiveness of rod bipolar cells due to a reduced rod input ^126^ while the sensitivity (basically the function of the cells) is stable during the intermediate phase (4 months). Finally, at late stage (6 months), rod-bipolar cells display a reduced responsiveness and a decline in sensitivity associated with retinal remodeling affecting all inner neurons as it occurs at late stages of disease ^30, 46–48^. The similarities between these disease studies and our model suggests that besides direct damage to photoreceptors or RPE, as it occurs in many inherited retinitis pigemntosa forms, impaired/malfunctioning MG are very likely contributing substantially to the overall disease progression. More importantly, we showed that these pathological events were not caused by MG absence ^19, 20^ but miRNA imbalances in the glia.

However, while cone impairments occurred at the same stage after Dicer loss, the rod loss was more accelerated in the Rlbp-Cre-driven strain which had a partially impaired RPE. The RPE has a variety of pivotal functions required for photoreceptor health and overall retinal health including continuous renewal of the photoreceptor outer segments, isomerization of all-trans-retinal to 11-cis-retinal and the formation of the blood-retinal barrier ^127–129^. Because of all these vital functions, impairments of the RPE does not only accompany but can lead to photoreceptor degeneration ^3, 128, 130, 131^.

Since miRNAs play a significant role in regulating essential biological processes by targeting networks of functionally correlated genes, they are vital components of the molecular networks in the RPE ^132, 133^. In fact, RPE Dicer-cKO studies as well as studies on selected miRNAs showed that RPE miRNAs are required for proper RPE maturation and function ^6, 7, 10, 130, 134–139^.

In the Rlbp-Dicer-cKO, we found obvious changes Dicer/miRNA deprived RPE including disrupted expression of Zo.1, dislocation of RPE nuclei, and enlarged cell size. Altered Zo1 expression and enlarged cell size are a known pathophysiological feature indicating RPE malfunction ^140–143^. More importantly, drastically altered Zo.1 expression was reported previously after Dicer deletion ^7, 10^ as well as after DiGeorge Critical Receptor 8 (Dgcr8) deletion, confirming that not just Dicer loss *per se* but miRNA cause RPE alteration ^7^. Dgcr8 is a part of the Drosha complex that generates precursor miRNAs from primary miRNA transcripts. This study also showed that the observed alterations were due to miRNA loss and not due to Alu element accumulations which are known to contribute to photoreceptor loss in AMD ^7, 144^. Since Alu elements were also absent in the datasets obtained from FACS-purified MG from the Rlbp-CreER:Dicer-cKO mice, it is rather unlikely that they play a substantial role in the observed phenotypes ^9, 21^.

Together our data shows that dysregulated but physically still present MG lead to cone impairment in an otherwise healthy retina. This suggests an additional or even new prospect for the development of retinal diseases, in particular cone dystrophies of unknown origin. Furthermore, this shows that drastic molecular changes long precede histological and functional changes. Histological manifestation (clinical observation) occurs relatively late which might be the reason for difficulties in treating such complex diseases.

## Acknowledgements

The authors thank Monica Andrade for her assistance with this study. This study was funded by the New York Empire Innovation Program Grant to S.G.W, the National Eye Institute (NEI, R01 EY032532) to S.G.W., SUNY Startup Funds to S.G.W, SUNY Graduate Assistantship to D.L., S.K. and S.C., NEI T35 to A.M.R, NEI R01 EY024373 and R21 EY032724 to S.F. Catalyst Award from Research to Prevent Blindness Inc./American Macular Degeneration Foundation to S.F., an unrestricted award to the Department of Ophthalmology and Visual Sciences from Research to Prevent Blindness, Inc.

## Notes

**Funding:** This study was funded by the New York Empire Innovation Program Grant to S.G.W, the National Eye Institute (NEI, R01 EY032532) to S.G.W., SUNY Startup Funds to S.G.W, SUNY Graduate Assistantship to D.L., S.K. and S.C., NEI T35 to A.M.R, NEI R01 EY024373 and R21 EY032724 to S.F. Catalyst Award from Research to Prevent Blindness Inc./American Macular Degeneration Foundation to S.F., an unrestricted award to the Department of Ophthalmology and Visual Sciences from Research to Prevent Blindness, Inc.

### Competing Interest Statement

The authors have declared no competing interest.

