## Supplementary Figures and Tables for "Dicer loss in Müller glia leads to a defined sequence of pathological events beginning with cone dysfunction"

### Supplementary material

Larbi et al.

#### Supplementary Table S1: Genotyping Primers

| Gene name | Forward sequence (5' to 3') | Reverse sequence (3' to 5') |
|---|---|---|
| **Rlbp1Cre transgene** | CAA GTG TGA GAG ACA GCA TTG | TCC TTA GCG CCG TAA ATC AA |
| **tdTomato wildtype** | AAG GGA GCT GCA GTG GAG TA | CCG AAA ATC TGT GGG AAG TC |
| **tdTomato mutant** | CTG TTC CTG TAC GGC ATG G | GGC ATT AAA GCA GCG TAT CC |
| **Dicer** | CCTGACAGTGACGGTCCAAAG | CATGACTCTTCAACTCAAACT |
| **Glast1Cre transgene** | ACA ATC TGG CCT GCT ACCAAA GC | CCA GTG AAA CAG CAT TGCTGT C |
| **RD 10** | GGC CAG TGA GAA CAA GGA AC | TGA TTC ATC TAG CCC ATC CA |

**Supplementary Table S2: Primary and Secondary antibodies**

| antibody | concentration | Company, Catalog # |
|---|---|---|
| **Primary antibodies** | | |
| rat anti RFP (tdTomato) | 1:1000 | Antibodies online, ABIN334653 |
| mouse anti glutamine synthetase (GS) | 1:500 | Millipore, MAB302 |
| rabbit anti Sox9 | 1:250 | Millipore, AB5535 |
| rabbit anti GFAP | 1:1000 | Dako, Z033401-2 |
| goat anti Otx2 | 1:250 - 1:500 | R&D Systems, AF1979 |
| rabbit anti Zo-1 | 1:300 | Invitrogen, 61-730-0 |
| rabbit anti Calretinin | 1:250 | Millipore, C7479 |
| mouse anti Calbindin | 1:500 | Millipore, ABN 2192 |
| goat anti Chat | 1:250 | Millipore, AB144P |
| rabbit anti M-opsin | 1:600 | Millipore, AB5405 |
| **Secondary antibodies** | | |
| Rhodamine Red 570 - AffiniPure F(ab')$_2$ Fragment Donkey Anti-Rat IgG (H+L) | 1:1000 | Jackson ImmunoResearch Laboratories, Inc. 712-296-150 |
| Alexa Fluor 488- AffiniPure F(ab')$_2$ Fragment Donkey Anti-Mouse IgG (H+L) | 1:500 | Jackson ImmunoResearch Laboratories, Inc. 715-546-150 |
| Alexa Fluor 488 - AffiniPure F(ab')$_2$ Fragment Donkey Anti-Rabbit IgG (H+L) | 1:500 | Jackson ImmunoResearch Laboratories, Inc. 711-546-152 |
| Alexa Fluor 488 - AffiniPure F(ab')$_2$ Fragment Donkey Anti-Goat IgG (H+L) | 1:500 | Jackson ImmunoResearch Laboratories, Inc 705-546-147 |
| Alexa Fluor 647 - AffiniPure F(ab')$_2$ Fragment Donkey Anti-Mouse IgG (H+L) | 1:500 | Jackson ImmunoResearch Laboratories, Inc 715-606-150 |
| Alexa Fluor 647 - AffiniPure F(ab')$_2$ Fragment Donkey Anti-Rabbit IgG (H+L) | 1:500 | Jackson ImmunoResearch Laboratories, Inc. 711-606-152 |
| Alexa Fluor 647 - AffiniPure F(ab')$_2$ Fragment Donkey Anti-Goat IgG (H+L) | 1:500 | Jackson ImmunoResearch Laboratories, Inc. 705-606-147 |

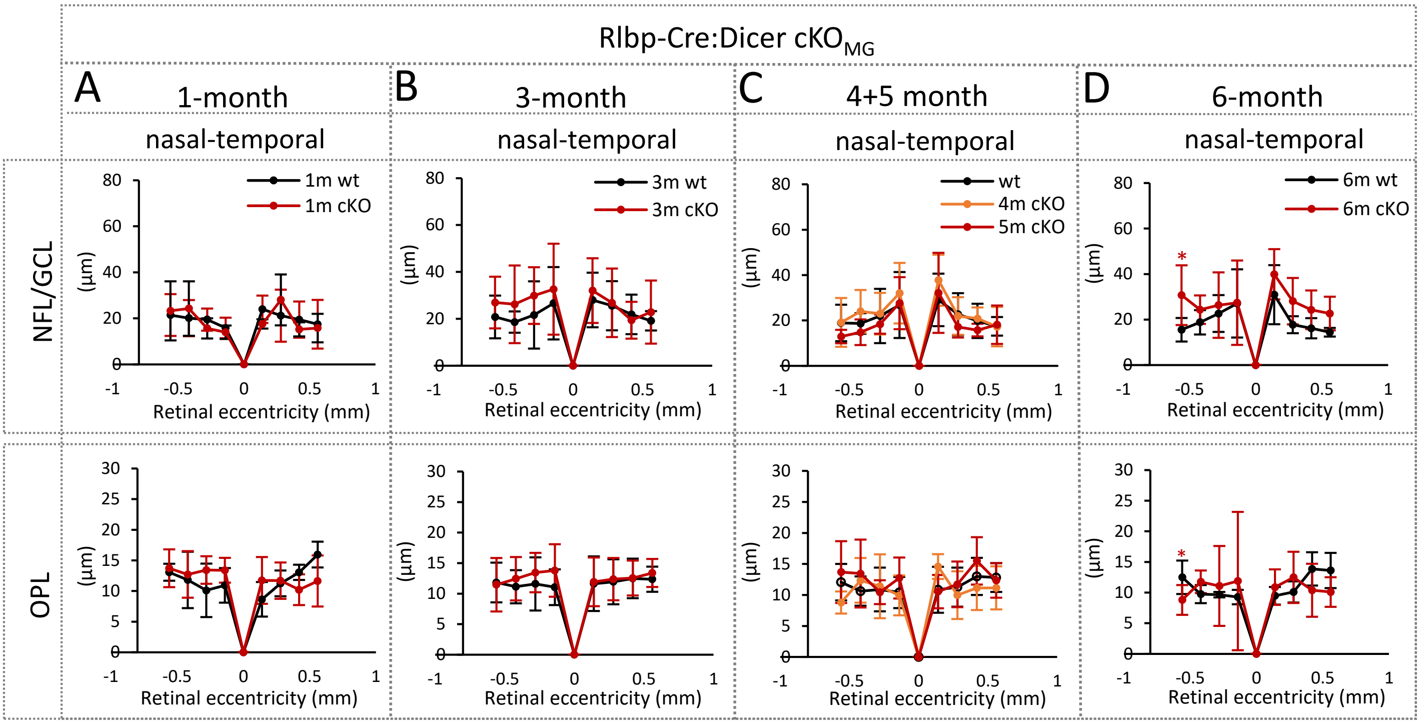


**Supplementary Figure S1: Müller glia alterations have no impact on the thickness of retinal ganglion cell layer and outer plexiform layer. A-D:** Spider plots of the thickness (µm) of the nerve fiber layer/ganglion cell layer (NFL/GCL) and outer plexiform layer (OPL) measured at the nasal-temporal axis of wildtype mice 1 month (n=4), 3 months (n=7), 4.5 months (n=11), or 6 months (n=4), and Rlbp-Cre:Dicer cKO$_{MG}$ mice (R-cKO$_{MG}$, cKO), 1 month (n=6), 3 months (n=11), 4 months (n=5), 5 months (n=5) or 6 months (n=7) after Cre induction. Mean ± S.D., significant differences are indicated, Mann-Whitney-U-test: *: $p \leq 0.05$. GCL: ganglion cell layer, IPL: inner plexiform layer, INL: inner nuclear layer, OPL: outer plexiform layer, ONL: outer nuclear layer, ELM: external limiting membrane.

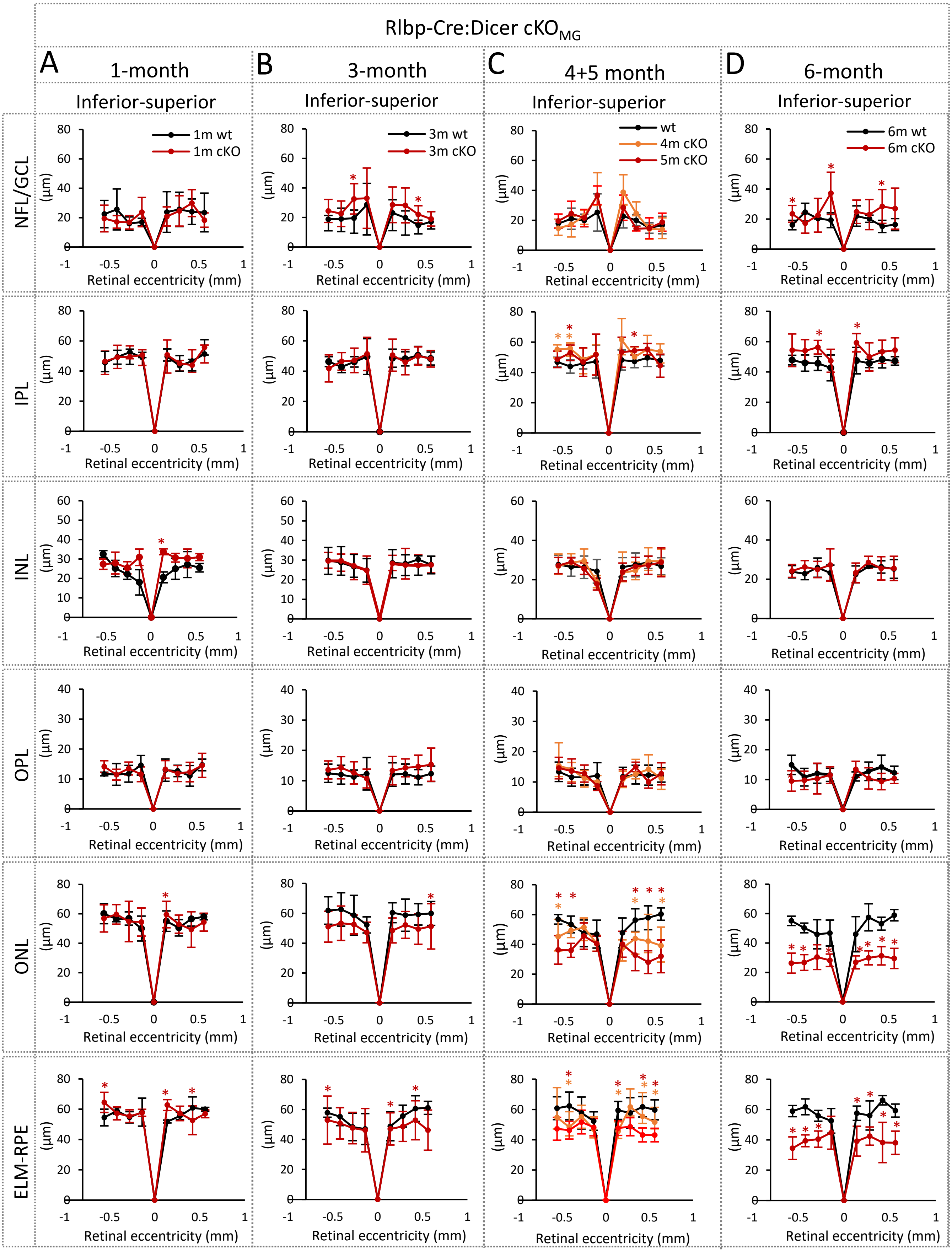


**Supplementary Figure S2: Müller glia alterations cause a reduction of retinal thickness in inferior-superior ELM/RPE.** Spider plots of the thickness (µm) of the nerve fiber layer/ganglion cell layer (NFL/GCL), inner plexiform layer (IPL), the inner nuclear layer (INL), outer plexiform layer (OPL), the outer nuclear layer (ONL) and external limiting membrane/retinal pigment epithelium (ELM-RPE) measured at the inferior-superior axis of wildtype mice 1 month (A, n=4), 3 months (B, n=7), 4+5 months (C, n=11), or 6 months (D, n=4), and Rlbp-Cre:Dicer cKO$_{MG}$ mice (cKO) 1 month (A, n=6), 3 months (B, n=11), 4+5 months (C, n=10), or 6 months (D, n=7) after Cre induction. Mean ± S.D., significant differences are indicated, Mann-Whitney-U-test: *: $p \leq 0.05$.

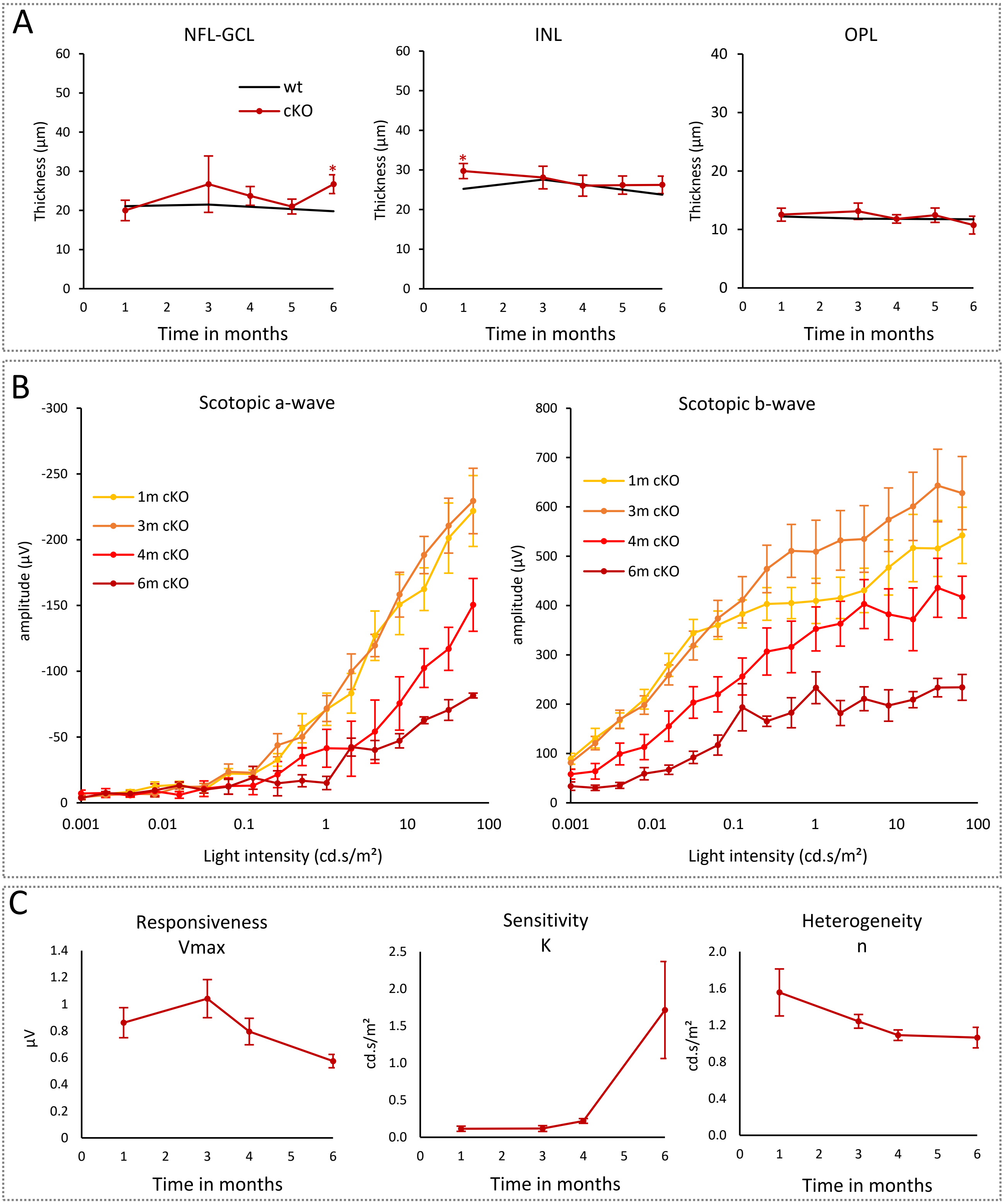


**Supplementary Figure S3: Rlbp-Cre:Dicer-cKO mice display progressive rod pathway loss over time. A**: Timeline plots showing averaged retinal layer thickness changes from OCT scans over time: wildtype 1 month (n=4), 3 months (n=7), 4 months (n=5), 5 months (n=5) and 6 months (n=4) and Rlbp-Cre:Dicer-cKO$_{MG}$, 1 month (n=6), 3 months (n=11), 4 months (n=5), 5 months (n=5) and 6 months (n=7). **B.** Full-field scotopic electroretinogram (ERG) recordings of the timeline showing a-wave and b-wave amplitudes of Rlbp-Cre:Dicer-cKO$_{MG}$ mice, 1 month (n=6), 3 months (n=8), 4 months (n=5), and 6 months (n=5). **C.** Timeline plots showing changes in the normalized saturated amplitudes (Vmax, responsiveness) semi-saturation (sensitivity), and slope (heterogeneity) using the Naka-Rushton equation over time. OCT: mean ± S.D., ERG: mean ± S.E.M. Significant differences are indicated, Mann-Whitney-U-test: *: $p \leq 0.05$.

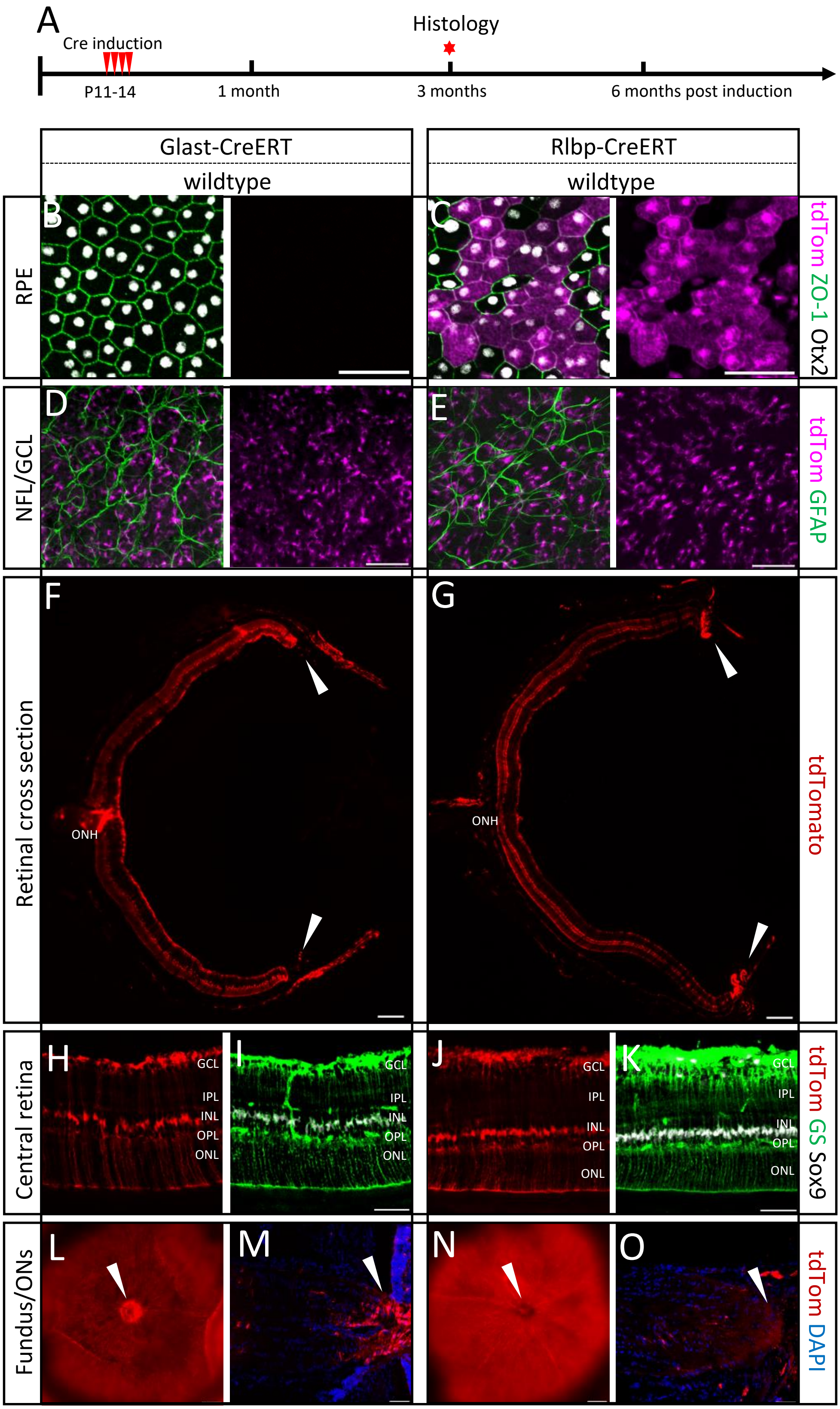


**Supplementary Figure S4: Glast-Cre reporter mice exclusively label Müller glia in the retina. A.** Experimental design. **B-M:** Endogenously labeled (tdTomato, tdTom) or immunofluorescence labeled cells of the retinal pigment epithelium (RPE, B-C), nerve fiber layer (NFL)/ganglion cell layer (GCL) of retinal flat mounts (D-E), whole retinas (F-G), central retinal areas of cross sections (H-K), or fundus (L, N) or optic nerve (ON) cross section (M, O) of Rlbp-Cre or Glast-Cre reporter mice using antibodies against RFP (tdTomato), ZO-1, Otx2, GFAP, Glutamine Synthetase (GS) and Sox9 as well as DAPI nuclear staining. Scale bars in B-E, H-J, N,O: 50 µm in F, G, L and M: 200 µm. Layer explanation is given in Figure 1.

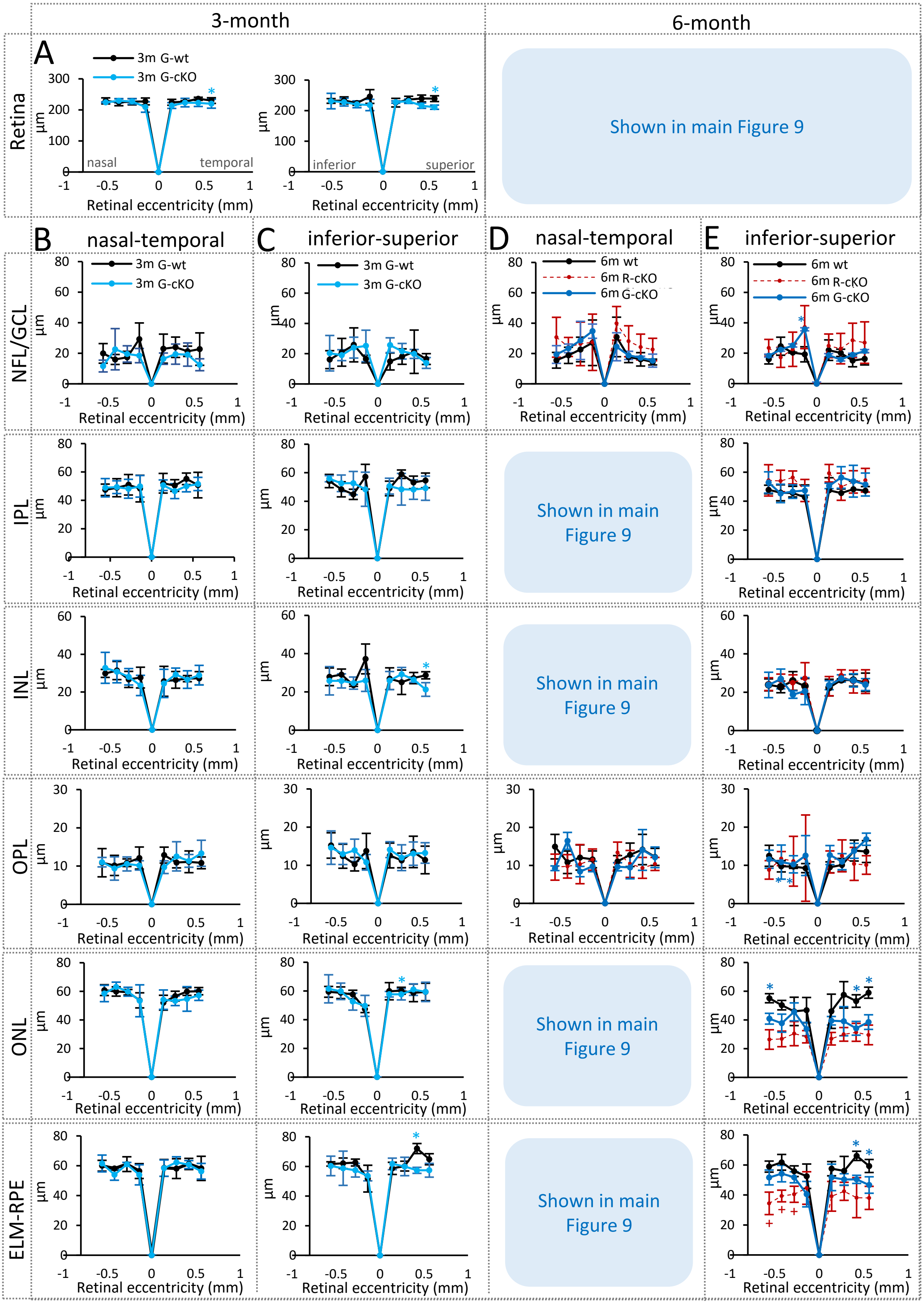


**Supplementary Figure S5: Glast-Cre:Dicer-cKO$_{MG}$ and Rlbp-Cre-Dicer cKO$_{MG}$ mice display similar reductions of retinal layers. A:** Spider plots of the overall retinal thickness (diameter, µm) measured at the nasal-temporal and superior-inferior axis of Glast-Cre wildtype (n=4) and Glast-Cre:Dicer-cKO$_{MG}$ mice (n=4). **B-E:** Spider plots of the thickness (µm) of the nerve fiber layer/ganglion cell layer (NFL/GCL), the inner plexiform layer (IPL), the inner nuclear layer (INL), the outer plexiform layer (OPL), the outer nuclear layer (ONL) and the external limiting membrane/retinal pigment epithelium (ELM-RPE), measured at the nasal-temporal or superior-inferior axis of 3-month Glast-Cre wildtype mice (G-wt, n=4), 3-month Glast-Cre:Dicer cKO$_{MG}$ mice (G-cKO, n=4, B-C), 6-month wildtype mice (n=4), 6-month G-cKO mice (n=6), 6-month Rlbp-Cre:Dicer cKO$_{MG}$ mice (R-cKO, n=7, D-E). Mean ± S.D., significant differences are indicated, (*): wt-cKO, (+) cKO-cKO comparisons, $p \leq 0.05$, Mann-Whitney-U-test.

Glast-CreER:Dicer-cKO$_{MG}$

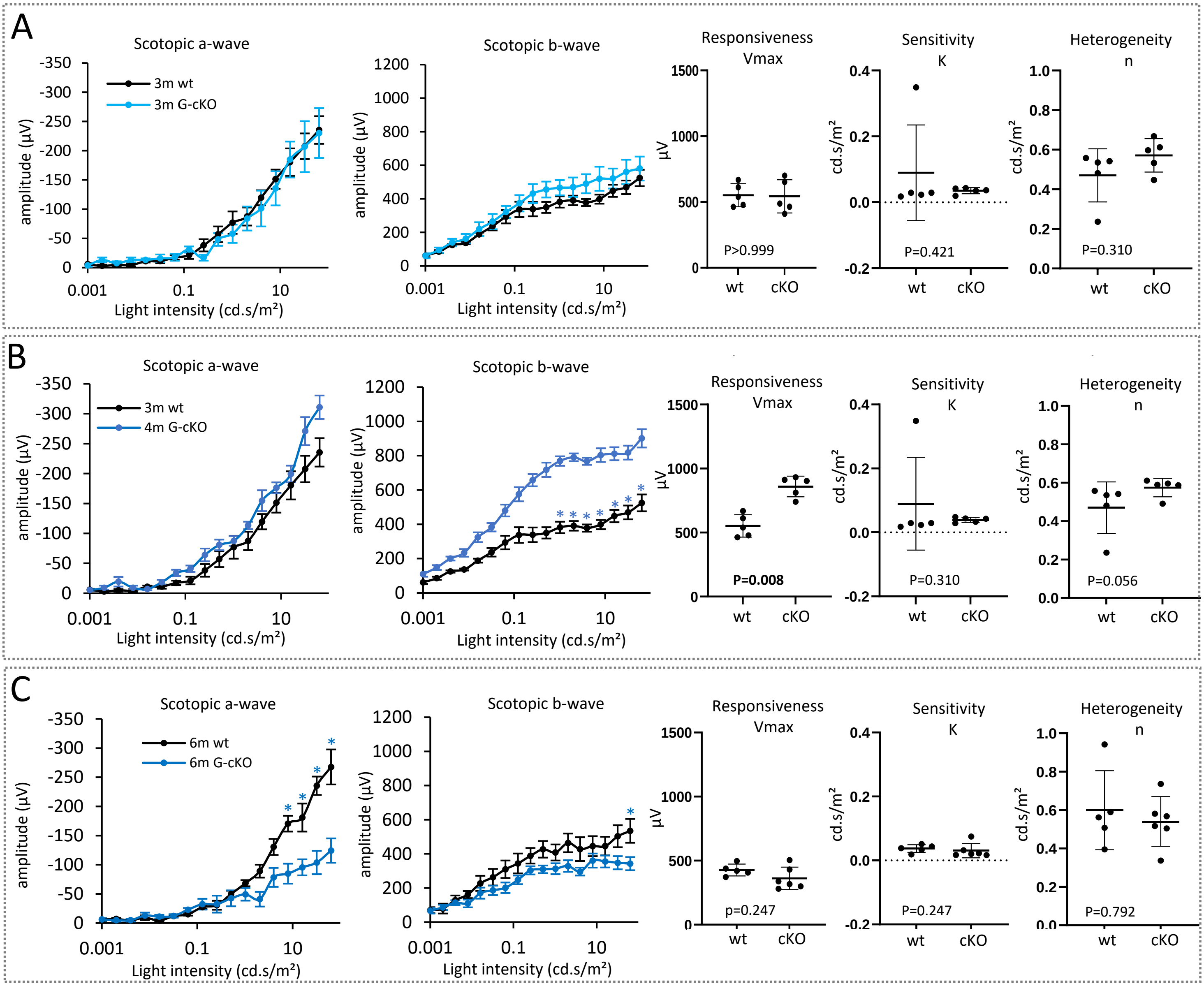


**Supplementary Figure S6: Glast-Cre:Dicer-cKO mice display first rod impairments 6 months after Cre induction.** Full-field scotopic electroretinogram (ERG) recordings showing a- and b-wave intensity-amplitude plots and estimated maximum amplitudes (Vmax, responsiveness), semi-saturation (sensitivity), and slope (heterogeneity) using the Naka-Rushton equation of 3-month Glast-Cre:Dicer-cKO$_{MG}$ mice (n=5) in comparison to wildtype (n=5, A), 4-month Glast-Cre:Dicer-cKO$_{MG}$ mice (n=5) in comparison to wildtype (n=5, B) and 6-month Glast-Cre:Dicer-cKO$_{MG}$ mice (n=5) in comparison to wildtype (n=5, C). Mean ± S.E.M., significant differences are indicated, Mann-Whitney-U-test: *: $p \leq 0.05$.
